# Complex opioid driven modulation of glutamatergic and cholinergic neurotransmission in a GABAergic brain nucleus associated with emotion, reward and addiction

**DOI:** 10.1101/2024.12.10.627344

**Authors:** R. Chittajallu, A. Vlachos, A.P. Caccavano, X.Q. Yuan, S. Hunt, D. Abebe, E. London, KA. Pelkey, C.J. McBain

## Abstract

The medial habenula (mHb)/interpeduncular nucleus (IPN) circuitry is resident to divergent molecular, neurochemical and cellular components which, in concert, perform computations to drive emotion, reward and addiction behaviors. Although housing one of the most prominent mu opioid receptor (mOR) expression levels in the brain, remarkably little is known as to how they impact mHb/IPN circuit function at the granular level. In this study, our systematic functional and pharmacogenetic analyses demonstrate that mOR activation attenuates glutamatergic signaling whilst producing an opposing potentiation of glutamatergic/cholinergic co-transmission mediated by mHb substance P and cholinergic neurons, respectively. Intriguingly, this latter non-canonical augmentation is developmentally regulated only emerging during later postnatal stages. In addition, we reveal that specific potassium channels act as a molecular brake on nicotinic receptor signaling in the IPN with the opioid mediated potentiation of this arm of neurotransmission being operational only following attenuation of Kv1 function. Thus, mORs play a complex role in shaping the salience of distinct afferent inputs and transmitter modalities that ultimately influences synaptic recruitment of downstream GABAergic IPN neurons. Together, these observations provide a framework for future investigations aimed at identifying the neural underpinnings of maladaptive behaviors that can emerge when opioids, including potent synthetic analogs such as fentanyl, modulate or hijack this circuitry during the vulnerable stages of adolescence and in adulthood.

## INTRODUCTION

Substance use disorders (SUDs) are agnostic to demographic group posing a widespread public health crisis impacting many communities^1^. Drugs of misuse influence synaptic function resulting in acute and protracted circuit adaptations that can exacerbate compulsive behaviors resulting in dependence^2,3^. Mu-opioid receptors (mORs) represent the primary molecular substrate responsible for both the clinically efficacious (i.e. pain management) and the euphoric effects promoting opioid misuse and addiction^4^. Additionally, mainly via liberation of endogenous opioids^5^, mORs also facilitate the addictive propensity of other substances^5–7^ e.g. alcohol and nicotine which, together with opioids, comprise the three most prevalent drugs of abuse worldwide^1^. mORs are expressed by varied neural cell-types resident in numerous regions within the brain^7–9^. Thus, detailed investigations as to how mOR activation impacts circuit dynamics constitutes an important endeavor critical for identification of potential therapeutic interventions aimed at alleviating the deleterious societal consequences of SUDs.

Here, we focus on a relatively understudied brain circuit comprising the medial habenula (mHb) and interpeduncular nucleus (IPN). The mHb is a bilateral epithalamic structure receiving significant synaptic input from, for example, the diagonal band and various septal regions^10–12^.

The efferent output of the mHb is predominantly mediated by two distinct populations, substance P (SP) and cholinergic neurons, that primarily impinge on the IPN^11,13^. This latter structure in the ventral midbrain is mainly populated with GABAergic neurons that influence activity in downstream brain regions such as the raphe nuclei and ventral tegmental area^14^. Thus, the mHb/IPN axis is anatomically positioned to integrate incoming limbic forebrain signals to ultimately control the function of these midbrain monoaminergic nuclei. It is therefore unsurprising that, perturbation/modulation of the synaptic recruitment or activity of IPN GABAergic neurons precipitates varied emotion, reward and addiction phenotypes^13–22^.

The mHb/IPN axes houses one of the highest densities of mORs in the central nervous system^23,24^ thus representing a prominent brain locus for opioid action. Despite this, descriptions regarding the role of mHb/IPN mORs in shaping behavior are still in their relative infancy^25,26^. mOR activation and subsequent G-protein signaling (Gi/o) classically produces an acute direct inhibitory effect in many neuronal types typified by membrane potential hyperpolarization via activation of inward rectifying potassium (GIRK) channels and/or reduced function of voltage-gated calcium channels function that are coupled to neurotransmitter release^8,27^. Strikingly, there is a dearth of information concerning the cellular effects and subsequent circuit influence of mOR signaling in the mHb/IPN rendering the neural correlates underlying the behavioral outcomes elicited by opioids unclear at present. Interestingly, direct excitatory effects of mOR activation are apparent in certain areas of the nervous system either under basal conditions or following chronic drug regimens^28–31^. Furthermore, within the IPN itself, GABA_B_-receptor activation (another Gi-linked receptor) produces an increase in synaptic transmission^32–34^ questioning whether this surprising effect is generalized to other members of this receptor family expressed within this region such as mORs.

The central role of the mHb/IPN circuitry underlying numerous behavioral aspects relating to emotion and the cycle of addiction, the conspicuous expression of mORs and a clear precedent for non-canonical effects of Gi-linked receptors together formed the impetus for the current work. Employing cell-type conditional optogenetics in combination with slice electrophysiology and pharmacogenetics we systematically dissect the effects of opioids on the mHb/IPN circuitry with a particular focus on synaptic signaling between these structures.

## RESULTS

The two major neuronal populations of the mHb, substance P (SP) and cholinergic subtypes can be distinguished by virtue of *Tac1* and *ChAT* gene expression. Leveraging the Allen Brain Institute’s publicly available whole mouse brain 10X single cell RNAseq dataset (https://alleninstitute.github.io/abc_atlas_access/intro.html)^35^ it is evident that these two neuronal classes within the mHb cluster (Subclass 145 MH) can be parsed based on their transcriptomic profiles (**Figure 1a**). Spatially, the SP and cholinergic neurons largely segregate to the dorsal versus ventral mHb, respectively **(Figure 1b,c)**. The projection pattern as delineated by use of specific Cre transgenic mouse lines (i.e ChATCre and Tac1Cre) when crossed to a conditional TdTomato reporter (Ai9), reveals a predominant innervation of IPL by SP neurons whereas cholinergic neurons impinge on the IPR/IPC subdvisions (**Figure 1c**).

**Figure 1.**
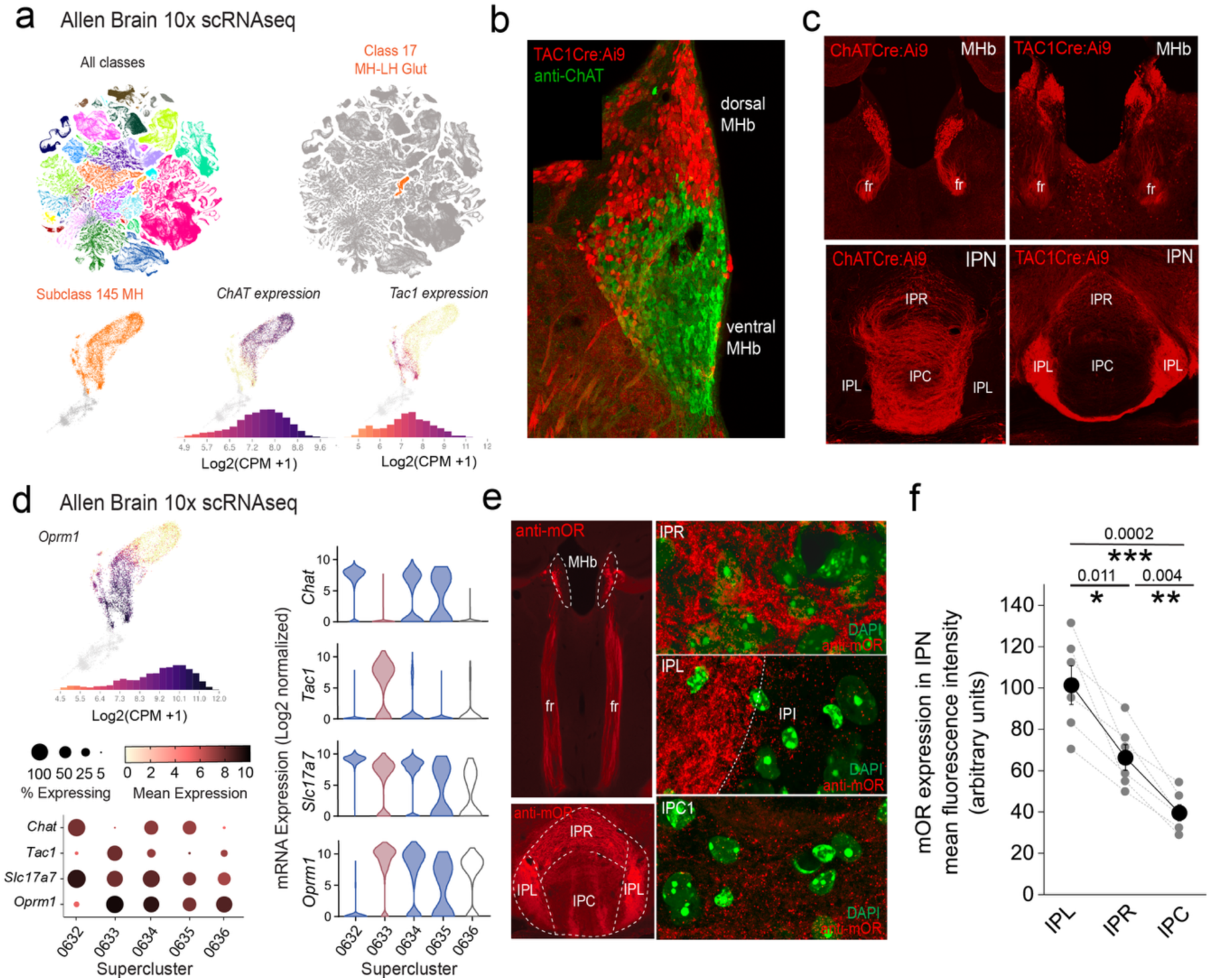
*OPRM1* gene and mOR protein expression in the habenula-interpeduncular axis. **(a)** 10X scRNAseq UMAP whole mouse brain showing all cellular classes (top left panel; 4.04 million cells) and Class 17 MH-LH Glut (right panel; corresponding to the mHB and lateral habenula; 10.8K cells). Bottom panels focus on individual mHB cells (Subclass 145 MH; 8K cells) indicating log2 mRNA expression of *ChAT* and *TAC1*. **(b)** Confocal image of a Tac1Cre:Ai9 mouse immunostained with anti-ChAT illustrating the distinct distribution of substance P (red) and cholinergic neurons (green) in dorsal and ventral mHb, respectively. **(c)** Conditional td-Tomato expression in cholinergic (left panels; ChATCre:Ai9) and SP neurons (right panels; Tac1Cre:Ai9) illustrating their spatial location within the mHB with their axonal outputs in the fasciculus retroflexus (fr) and largely non-overlapping terminal axonal arborization patterns in IPN. **(d)** Profile of log2 mRNA expression of *OPRM1* in Subclass 145 MH (top left panel). Corresponding dot and violin plots depicting 3 CHAT (0632,0634,0635) and 1 TAC1 (0633) cluster with relative expression of *Slc17a7* (VGluT1) and *OPRM1* in each corresponding cluster. In the violin plots the CHAT clusters and TAC1 superclusters are color coded blue and red, respectively. **(e)** Endogenous mOR expression throughout the mHb and IPN axis (left panels; red) assessed by immunocytochemistry. High resolution airy scan images of mOR distribution in subdivisions of the IPN; rostral IPN (IPR), lateral IPN (IPC) and central IPN (IPC). Green fluorescence are DAPI stained nuclei. (**f**) Densitometry analyses of mOR expression in the various subfields of IPN. Data are from 2-4 slices containing IPN taken from each of 6 mice aged P40-P60. Data depicted in **(a)** and **(d)** are from the publicly available Allen Brain cell Atlas (https://knowledge.brain-map.org/abcatlas). See methods for further details.

Together these distinct neuronal classes provide much of the afferent input to the IPN via the fasciculus retroflexus (fr) axonal tracts **(Figure 1c**) ultimately dictating IPN neuronal recruitment. At the mRNA level, two of the designated three ChAT neuronal superclusters and the single Tac1 supercluster demonstrate significant expression of *OPRM1* (mean expression levels including zero values = 0.33, 6.8 and 4.1 in *ChAT* Superclusters 0632, 0634, 0635, respectively and 8.7 in the *Tac1* Supercluster 0633; **Figure1d**) At the protein level, and as previously reported^23^, we confirm that mOR expression is notable in the mHb, the fasciculus retroflexus axonal tracts and particularly prevalent in the IPN **(Figure 1e)**. Within this latter structure, high resolution microscopy clearly show mOR expression is densest in the lateral IPN (IPL) with intermediate and relatively lower levels found in the rostral IPN (IPR) and central IPN (IPC) subregions, respectively (**Figure 1e,f**).

Selective stimulation of each of these mHb neuronal populations in isolation is achieved using Cre-mediated conditional expression of channel rhodopsin by crossing Ai32 mice with either ChATCre or TAC1Cre mice; see methods for details). Adopting this optogenetic approach we investigate how mOR receptor activation modulates habenulo-interpeduncular synaptic dialogue imparted by these parallel yet distinct afferent systems in adult mice (p>40). Both SP and cholinergic neurons express the glutamate vesicular transporter, VGluT1 (*Slc17a7*; **Figure1d**) and competently release this excitatory neurotransmitter^21,36,37^ .

### mOR activation reduces substance P neuronal mediated glutamatergic transmission onto IPL GABAergic neurons

Since the highest expression of mOR was found in the IPL (**Figure 1e,f**), we initially probed the effect of a selective mOR agonist (DAMGO, 500 nM) on AMPA receptor mediated transmission mediated by SP neurons that prominently innervate this subregion (**Figure 1c**). DAMGO application results in a significant decrease in light-driven (470 nM; typically, 10-50 % arbitrary LED power equating to approximately 0.4 – 3.4 mW/mm^2^; CoolLED illumination system) AMPAR mediated EPSCs in TAC1Cre:Ai32 mice that is partially reversed upon washout of the agonist (**Figure 2a,b**). This is accompanied by an increase in S2/S1 paired pulse ratio (PPR) suggesting the modulation is driven by presynaptic changes in release probability (**Figure 2c**). Thus, neurotransmitter release by this specific afferent input to the IPL is directly, negatively modulated like that seen in many other neural circuits within the brain^8^.

**Figure 2.**
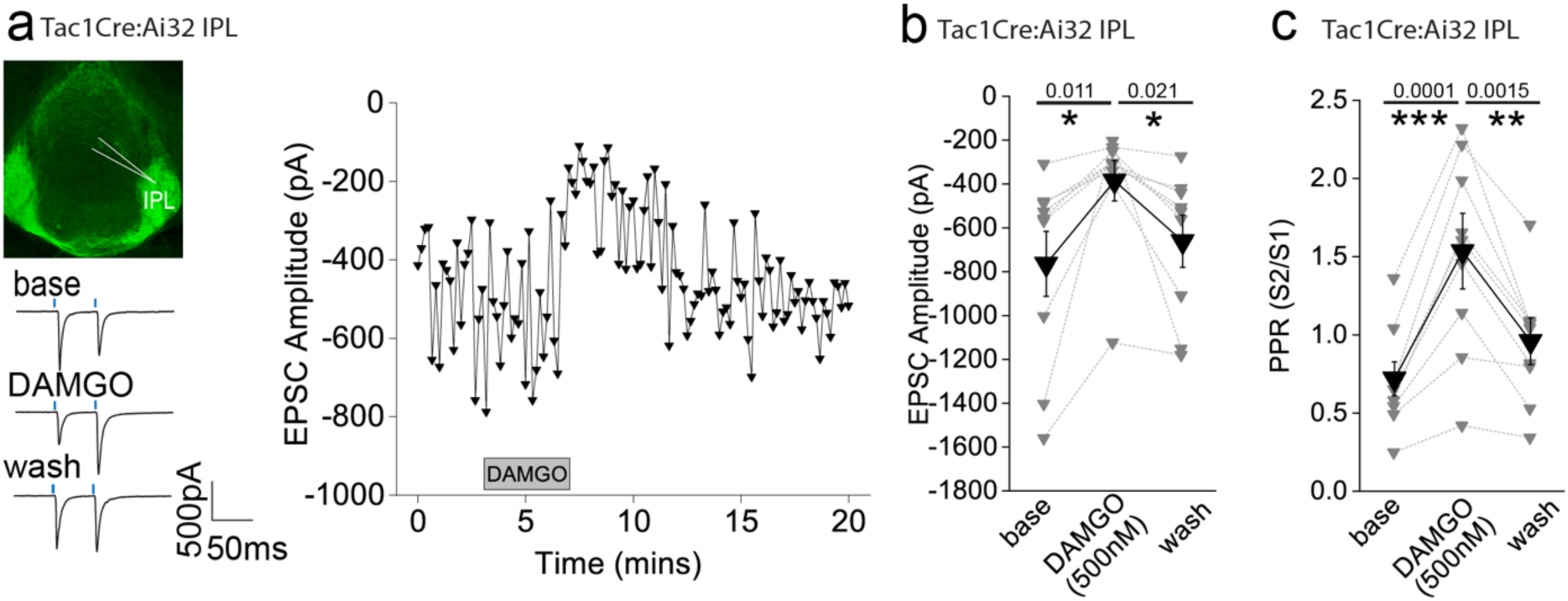
AMPAR-mediated synaptic transmission in lateral IPN (IPL) mediated by substance P neurons is inhibited by mOR activation(a) Whole-cell voltage-clamp in adult (>p40) Tac1Cre:Ai32 mice (top left panel illustrating the axonal arborization of ChR2 expressing SP neuronal axons in IPN and the position of neuronal recording in IPL). Single voltage-clamp traces of light evoked AMPAR EPSCs (bottom left panel; 470nM light pulse; 2 stimulations at 20Hz, blue dashes)) and time course of peak amplitudes (right panel) under baseline, during application of 500 nM DAMGO and washout conditions. **(b,c)** Individual (grey filled symbols) and mean (black filled symbols) data of AMPAR EPSC amplitude and corresponding paired pulse ratios (PPR; n=9 recorded IPL neurons from 7 mice).

### mOR activation augments the strength and fidelity of glutamatergic transmission mediated by cholinergic neurons resulting in enhanced excitation-spike coupling in IPR GABAergic neurons

We next examined how mORs modulate mHb-IPN neurotransmission that occurs via the other major input mediated by cholinergic neurons. To this end, we employed ChATCre:Ai32 mice and focused on the IPR (**Figure 3a**) due to the relatively high levels of combined mOR expression and cholinergic neuronal innervation (**Figure 1c,e,f**) in this subregion. Under our basal experimental conditions and in agreement with previous studies^21,37,38^ light evoked postsynaptic responses in the IPN mediated by mHb cholinergic neurons in response to either brief single or paired-pulse stimulation are solely mediated by AMPARs as evidenced by complete pharmacological block by DNQX (refer to **Figure 5-figure supplement 1a** and **Figure 7-figure supplement 1a-c**). Remarkably, DAMGO application elicited a significant robust and reversible potentiation of light-evoked (470 nM; typically, 10-100 % arbitrary LED power equating to approximately 0.4 – 6.9 mW/mm^2^; CoolLED illumination system) AMPAR EPSC amplitude (**Figure 3a,b**) in stark contrast to that seen with mHb TAC1-IPL glutamatergic signaling described previously (**Figure 2a,b**). Notably, the accompanying PPR changes were not consistent with an increase in release probability (**Figure 3c**)raising uncertainty as to the synaptic locus of the mOR effect.

**Figure 3.**
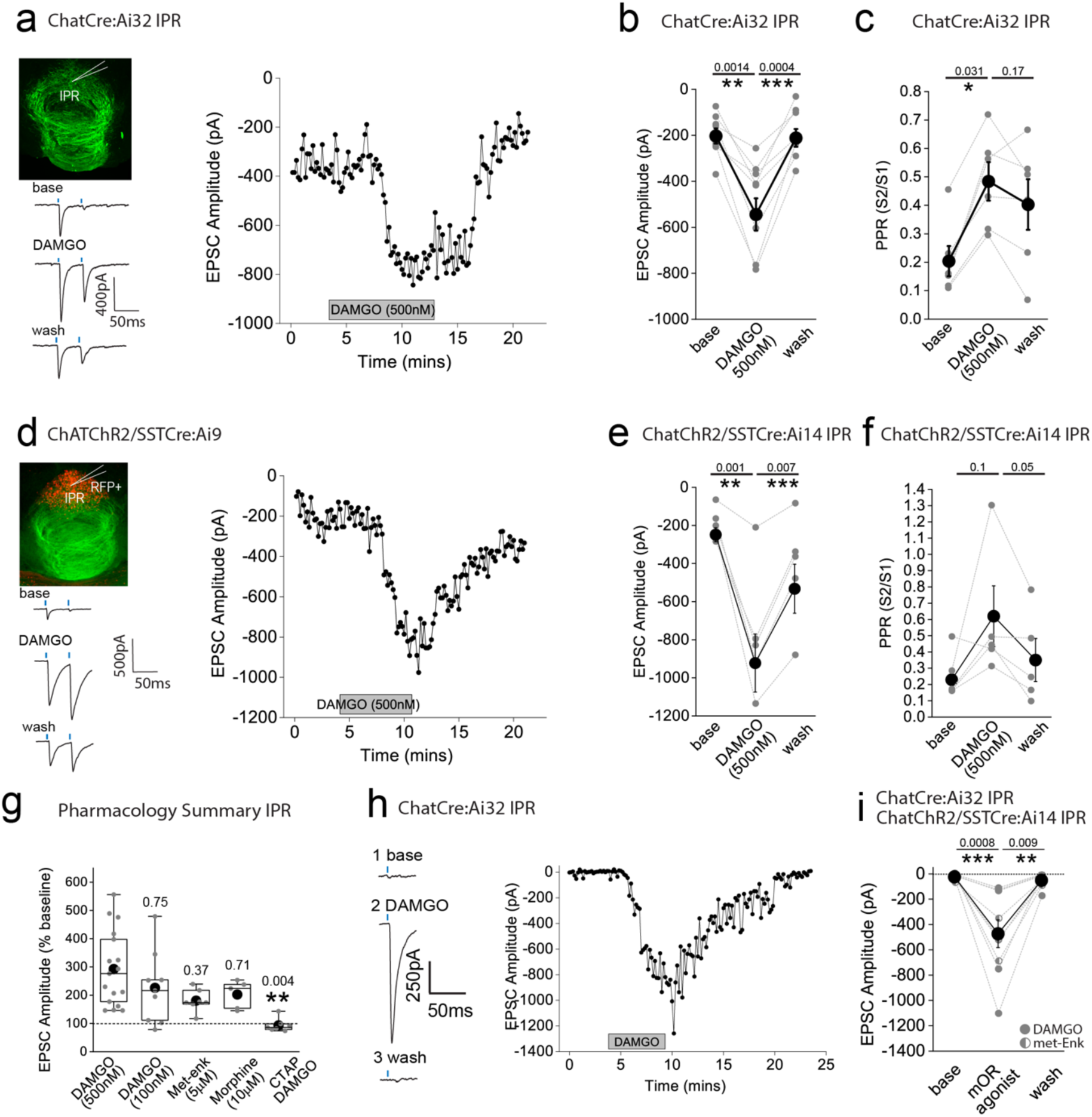
AMPAR-mediated synaptic transmission in rostral IPN (IPR) mediated by mHB cholinergic neurons is potentiated by mOR activation. **(a)** Whole-cell voltage-clamp in adult (>p40) ChATCre:Ai32 mice (top left panel illustrating the axonal arborization of ChR2 expressing cholinergic neuronal axons in IPN and the position of neuronal recording in IPR). Single voltage-clamp traces of light evoked AMPAR EPSCs (bottom left panel; 470nM light pulse; 2 stimulations at 20Hz, blue dashes) and time course of peak amplitudes (right panel) under baseline, during application of 500 nM DAMGO and washout conditions. **(b,c)** Individual (grey filled symbols) and mean (black filled symbols) data of AMPAR EPSC amplitude and corresponding paired pulse ratios (PPR; n=8 and 6 recorded IPR neurons from 7 and 5 mice for AMPAR EPSC amplitude and PPR, respectively). (**d**) Whole-cell voltage-clamp in adult (>p40) ChATChR2:SSTCre:Ai9 mice (top left panel illustrating the axonal arborization of ChR2 expressing cholinergic neuronal axons in IPN and the position of neuronal recording in RFP+ SST IPR neurons). Single voltage-clamp traces of light evoked AMPAR EPSCs (bottom left panel; 470nM light pulse; 2 stimulations at 20Hz, blue dashes) and time course of peak amplitudes (right panel) under baseline, during application of 500 nM DAMGO and washout conditions. **(e,f)** Individual (grey filled symbols) and mean (black filled symbols) data of AMPAR EPSC amplitude and corresponding paired pulse ratios (PPR; n=5 recorded IPR neurons from 4 mice). **(g)** Summary bar graph illustrating the effect of DAMGO (500nM and 100nM; n=16 recorded IPR neurons from 14 mice and n=9 recorded IPR neurons from 4 mice, respectively), met-enkephalin (5μm; n=7 recorded IPR neurons from 5 mice), morphine (10μM; n=5 recorded IPR neurons from 5 mice) and 500nM DAMGO in presence of mOR antagonist 1mM CTAP (n=7 recorded IPR neurons from 2 mice). **(h)** Voltage-clamp trace examples (single light stimulus; blue bar; left panel) and time course (right panel) under baseline, DAMGO and washout conditions in a recorded IPR neuron displaying no measurable baseline AMPAR EPSC in response to maximal light evoked stimulation (470nm; 6.9mW/mm^2^). **(i)** Individual (gray filled and half-filled symbols for 500nM DAMGO or 5μM Met-enkephalin respectively) and pooled (black filled symbols) data of light-evoked AMPAR EPSC amplitude during DAMGO application and washout (n = 6 and 3 recorded IPR cells from 6 and 3 mice for DAMGO and met-enkephalin application, respectively).

**Figure 3-figure supplement 1.**
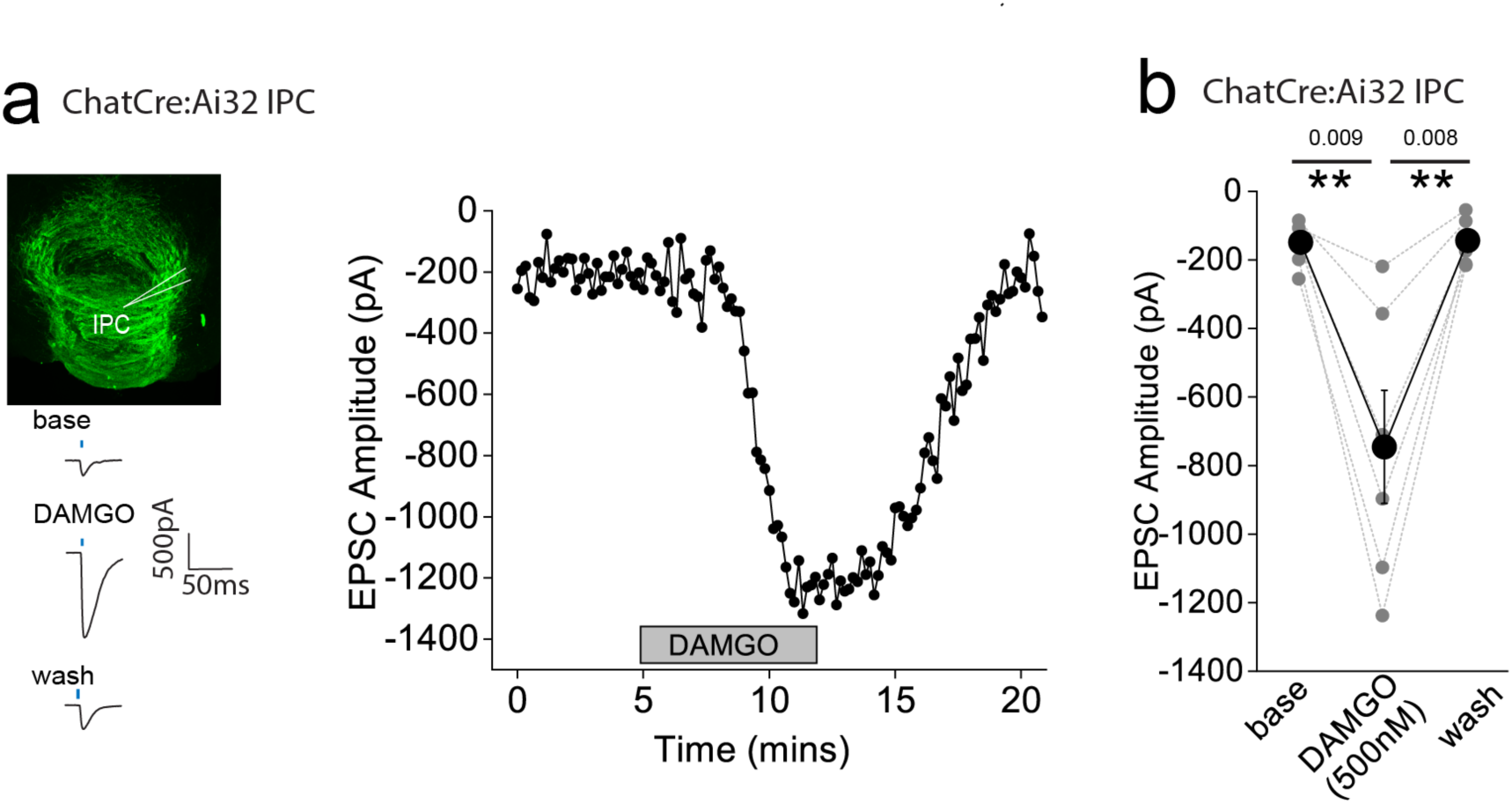
AMPAR-mediated synaptic transmission in central IPN (IPC) mediated by mHB cholinergic neurons is potentiated by mOR activation. **(a)** Whole-cell voltage-clamp in adult (>p40) ChATCre:Ai32 mice (top left panel illustrating the axonal arborization of ChR2 expressing cholinergic neuronal axons in IPN and the position of neuronal recording in IPC). Single voltage-clamp traces of light evoked AMPAR EPSCs (bottom left panel; 470nM light pulse; single stimulations, blue dash) and time course of peak amplitudes (right panel) under baseline, during application of 500 nM DAMGO and washout conditions. **(b)** Individual (grey filled symbols) and mean (black filled symbols) data of AMPAR EPSC amplitude (n=8 recorded IPC neurons from 7 mice).

**Figure 4.**
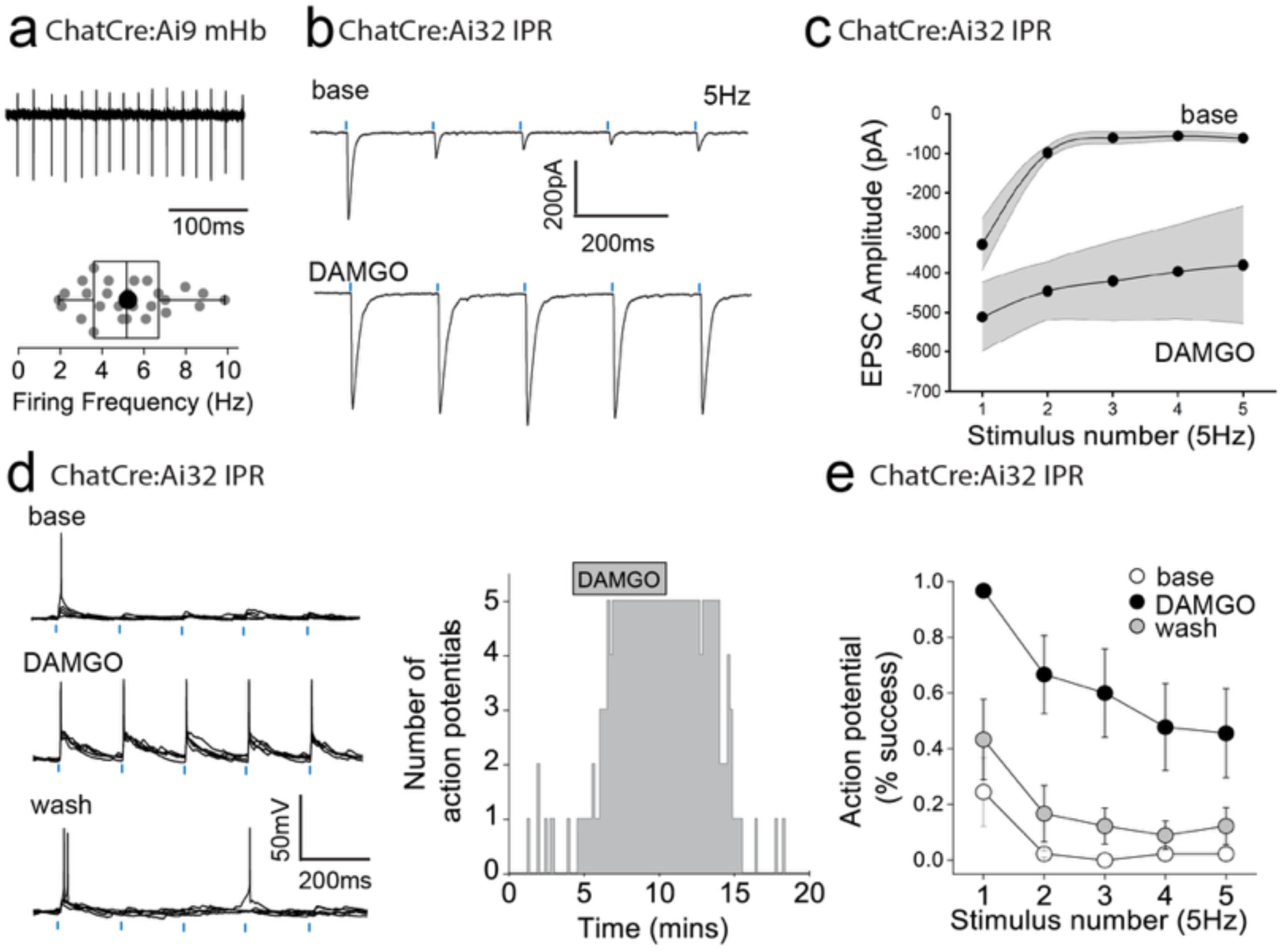
mOR activation increases fidelity of glutamatergic transmission mediated by mHb cholinergic neurons to augment excitation:spike coupling in postsynaptic IPR neurons. **(a)** Single example trace showing spontaneous action potential firing in cell attached mode from a td-Tomato-positive ventral mHb neuron in the ChatCre:Ai9 mouse (top panel). Box plot and corresponding individual data of the spontaneous firing frequency of mHB cholinergic neurons (bottom panel; n = 27 recorded cells). **(b)** Single voltage-clamp traces of light-evoked (470nm, 5 pulses delivered at 5Hz; blue dashes) AMPAR EPSCs recorded from postsynaptic IPR neurons in adult ChATCre:Ai32 mice (p>40) under baseline and following 500nM DAMGO application (left panel). **(c)** Pooled data (shaded area denotes SEM) of peak amplitude for each given stimulus in the 5Hz train during baseline and following 500nM DAMGO application (n = 5 recorded IPR neurons from 5 mice). **(d)** Single current-clamp traces example (5 consecutive overlaid sweeps) of light-evoked EPSP:action potential coupling (5 stimuli at 5 Hz; blue dashes) under baseline, 500nM DAMGO and washout conditions (left panel). Corresponding single example time course plot depicting number of light-driven EPSP-evoked action potential (right panel). **(e)** Mean data showing percentage success over 5 consecutive traces of EPSP:action potential coupling for each given stimulus in the 5Hz train under baseline (open symbols), 500nM DAMGO (black symbols) and washout (grey symbols) conditions (n = 9 recorded IPR neurons from 7 mice).

**Figure 5.**
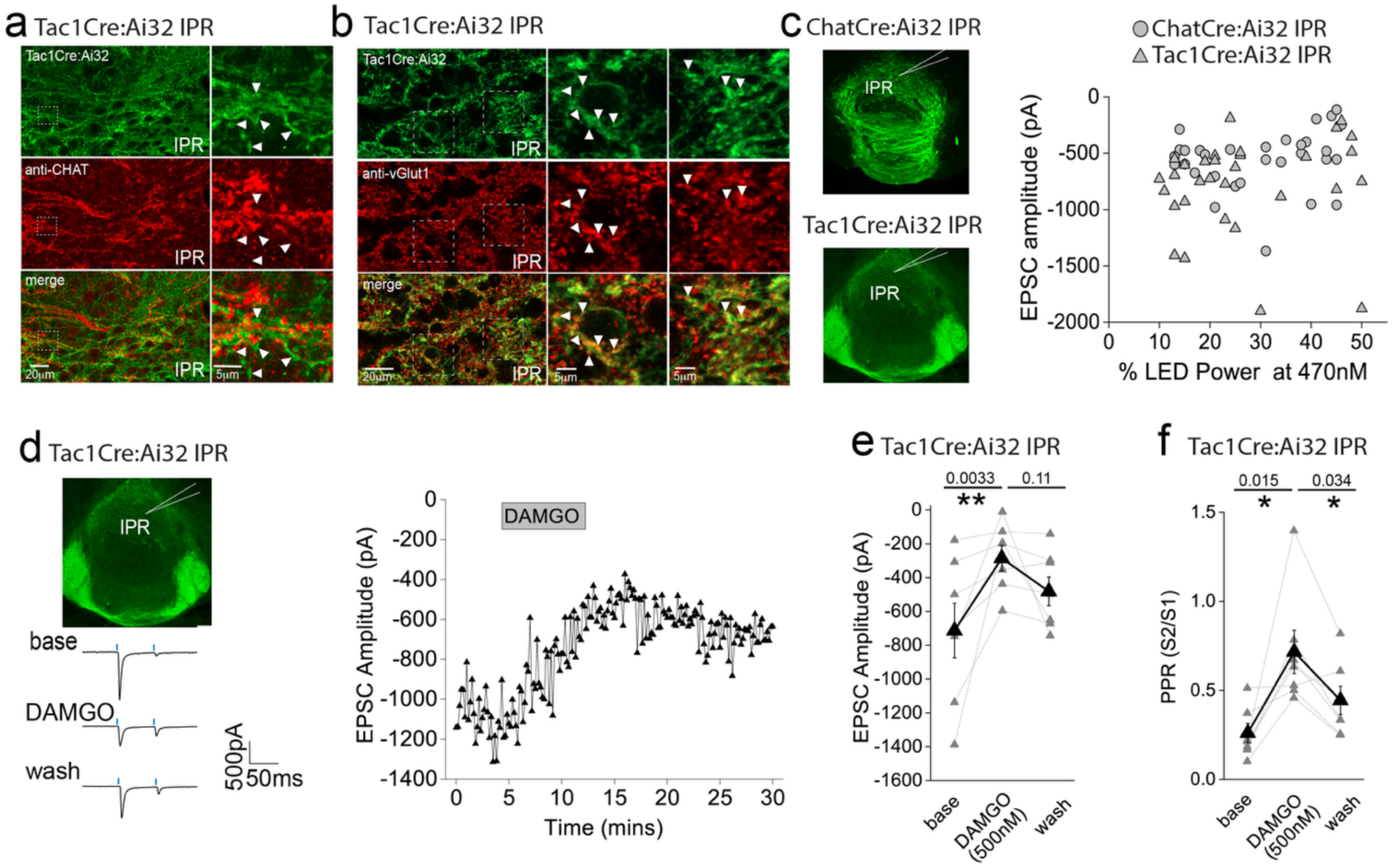
Effect of mOR activation on a novel functional SP neuronal mediated evoked AMPAR EPSCs in IPR. **(a)** High resolution airy scan of the IPR in TAC1Cre:Ai32 mouse showing ChR2-expressing synaptic bouton like-structures (green) and endogenous ChAT expression (red). Right panels are magnified regions of the boxed area. Arrows indicate examples of non-overlap of TacCre:Ai32 boutons with ChAT. **(b)** High resolution airy scan of the IPR in TAC1Cre:Ai32 mouse showing ChR2-expressing synaptic bouton like-structures (green) and endogenous VGlut1 expression via immunostaining (red). Right panels are magnified regions of the boxed area. Arrows indicate examples of expression of VGluT1 within TacCre:Ai32 bouton structures. **(c)** Comparison of light-evoked AMPAR EPSC peak amplitude in postsynaptic IPR neurons mediated by either cholinergic (ChATCre:Ai32 mice; n = 32 recorded IPR neurons) or substance P (Tac1Cre:Ai32 mice; n = 31 recorded IPR neurons) mHB neurons at various arbitrary % LED (10-50% corresponding to an approximate power of 0.4 – 3.4 mW/mm^2^; right panel). **(d)** Whole-cell voltage-clamp in adult (>p40) TAC1Cre:Ai32 mice (top left panel illustrating the axonal arborization of ChR2 expressing SP neuronal axons in IPN and the position of neuronal recording in IPR). Single voltage-clamp traces of light evoked AMPAR EPSCs (bottom left panel; 470nM light pulse; 2 stimulations at 20Hz, blue dashes) and time course of peak amplitudes (right panel) under baseline, during application of 500 nM DAMGO and washout conditions (right panel). **(e,f)** Individual (grey filled symbols) and mean (black filled symbols) data of AMPAR EPSC amplitude and corresponding paired pulse ratios (PPR; n=8 recorded IPR neurons from 7 mice).

**Figure 5-figure supplement 1.**
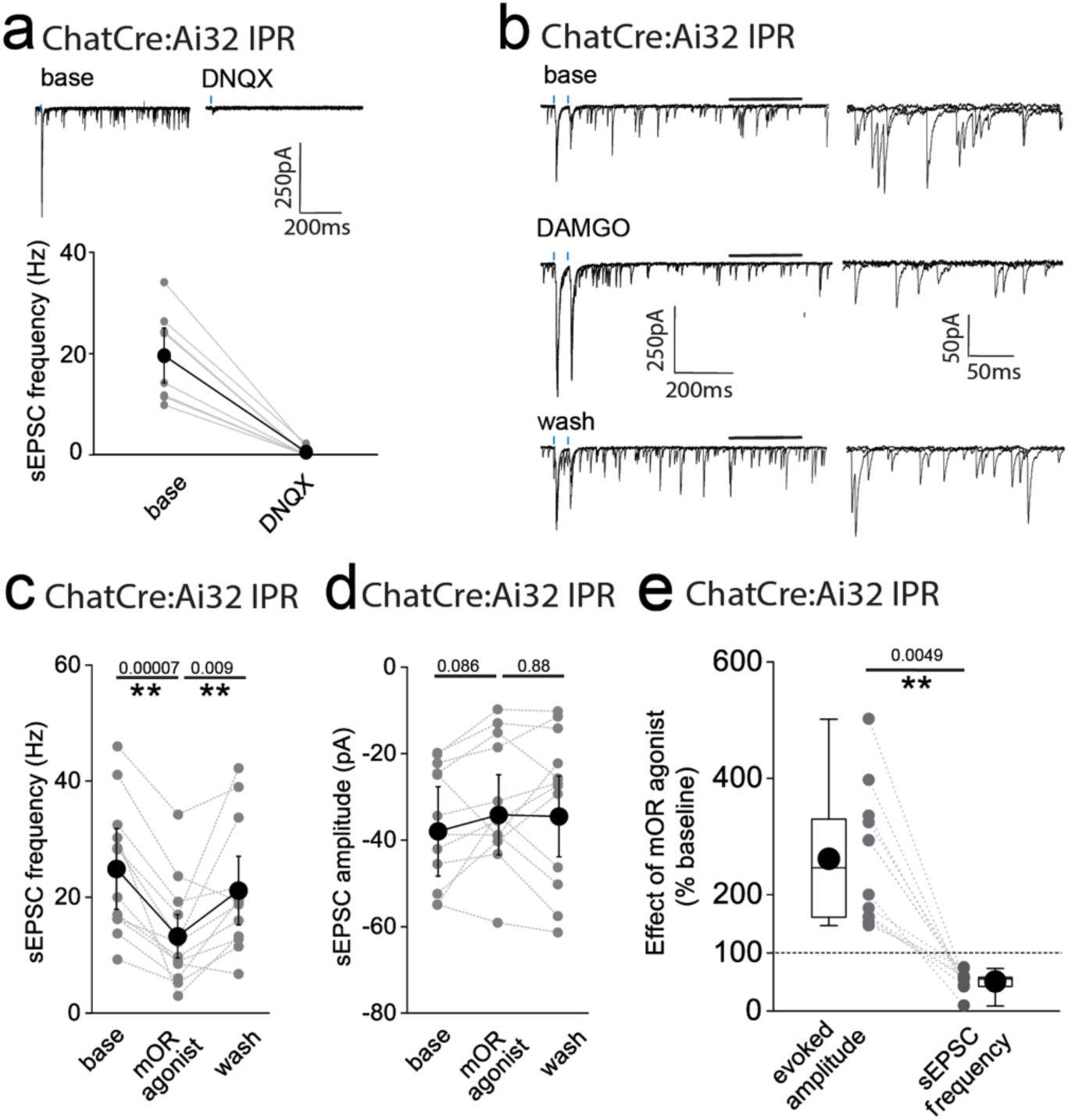
mOR activation imparts opposing effects on spontaneous AMPA EPSCs versus cholinergic neuron mediated evoked AMPAR EPSCs in IPR. **(a)** Single voltage-clamp traces (top panels) illustrating light-evoked AMPAR EPSCs mediated by cholinergic neurons and spontaneous EPSCs (sEPSCs) under baseline and following 10μM DNQX application in an IPR neuron. Individual (grey symbols) and pooled data (black symbols) illustrating complete cessation of sEPSCs as assessed by frequency following 10μM DNQX application (n = 8 recorded IPR neurons from 8 mice; bottom panel). **(b)** Single voltage-clamp traces under baseline, 500nM DAMGO and washout conditions of light-evoked AMPAR EPSCs mediated by mHb cholinergic neurons (2 stimuli at 20Hz; blue dashes) and sEPSCs (left panels). Magnification of sEPSCs events (right panels) corresponding to the region of the traces in the left panels delineated by the black bars. **(c,d)** Individual (grey symbols) and mean (black symbols) data of sEPSC frequency and amplitude (n = 12 recorded IPR cells from 11 mice). **(e)** Box plot and individual data depicting the relative DAMGO mediated percentage change from baseline of the light-evoked AMPAR EPSC peak amplitude versus AMPAR sEPSC frequency in each individual IPR neuron recorded (n = 12 recorded IPR cells from 11 mice).

**Figure 6.**
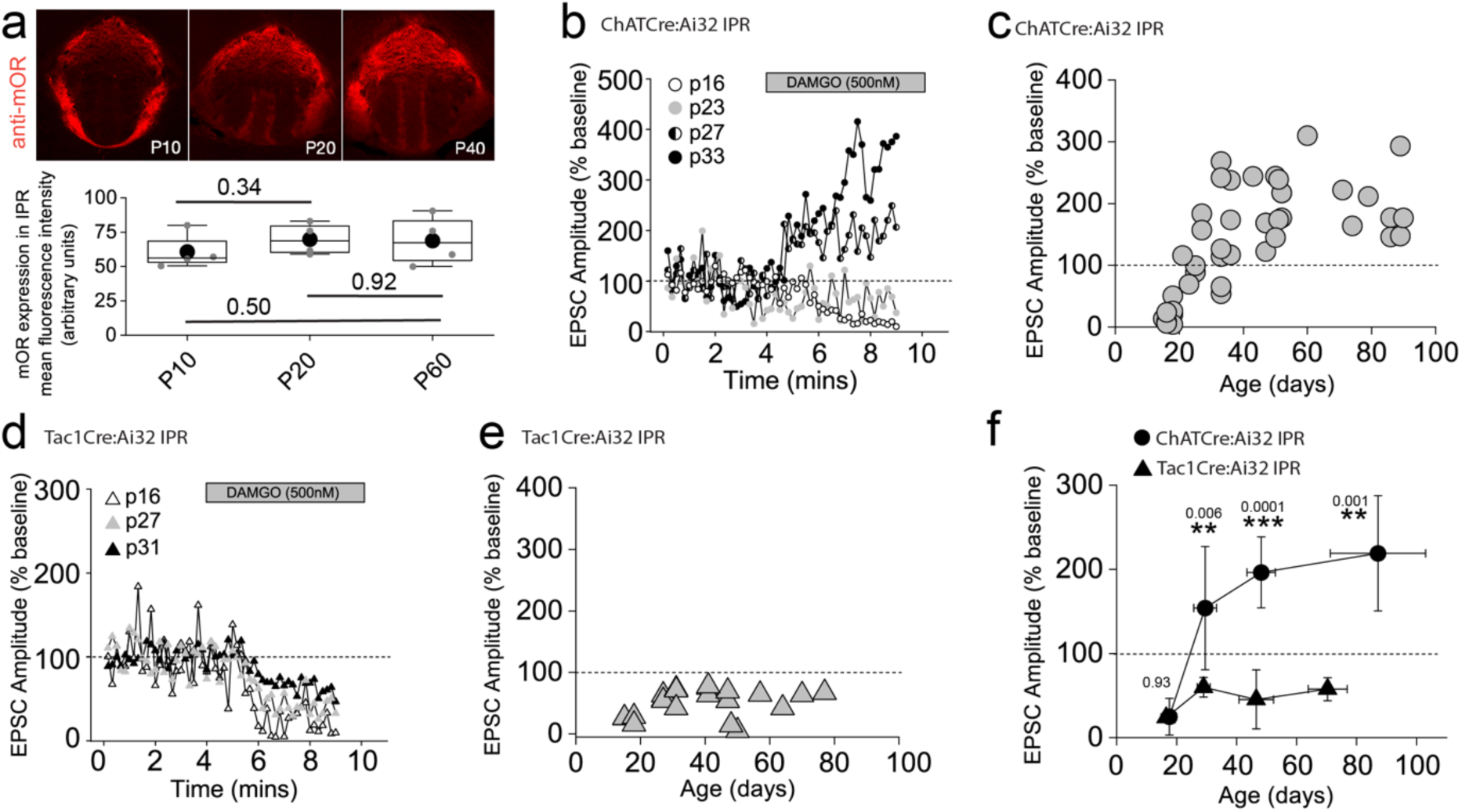
mORs constitute a developmentally regulated molecular switch altering the salience of neurotransmission in IPR mediated by substance P versus cholinergic neurons. **(a)** Confocal images of mOR protein expression in IPN during development (p10, p20 and p40). Densitometry analyses of mOR protein expression in IPR across development (measurements taken from 2 slices containing IPR from each of 2-4 mice for each age **(b)** Single examples of the time course of light-evoked AMPAR EPSC peak amplitudes mediated by mHB cholinergic neurons following DAMGO application in postsynaptic IPR neurons at varying ages as indicated. **(c)** Individual data of the percent change of light-evoked cholinergic neuronal mediated AMPAR EPSC peak amplitude elicited following DAMGO application across all ages tested (n = 44 recorded IPR neurons). **(d)** Single examples of the time course of light-evoked AMPAR EPSC peak amplitudes mediated by mHB substance P neurons following DAMGO application in postsynaptic IPR neurons at varying ages as indicated. **(e)** Individual data of the percent change of light-evoked SP neuronal mediated EPSC peak amplitude elicited following DAMGO application across all ages tested. **(f)** Summary plot of the mean changes in the normalized AMPAR EPSC peak amplitude (% baseline) by DAMGO mediated by cholinergic and SP neurons binned at the following developmental epochs; p15-23 (postnatal), p24-34 (adolescent/pre-pubescent), p35-60 (adolescent/pubescent, sexual maturation) and >p60 (adult)^96^. Total numbers of cells recorded = 39 and 21 cells from CHATCre:Ai32 and TAC1Cre:Ai32 mice, respectively. Error bars denote standard deviation of the mean. Note datapoints for ages >p40 in panels **c** and **e** are normalized data taken from recorded cells that were depicted in Figures 3b and **5e** as absolute peak amplitude changes, respectively.

**Figure 7.**
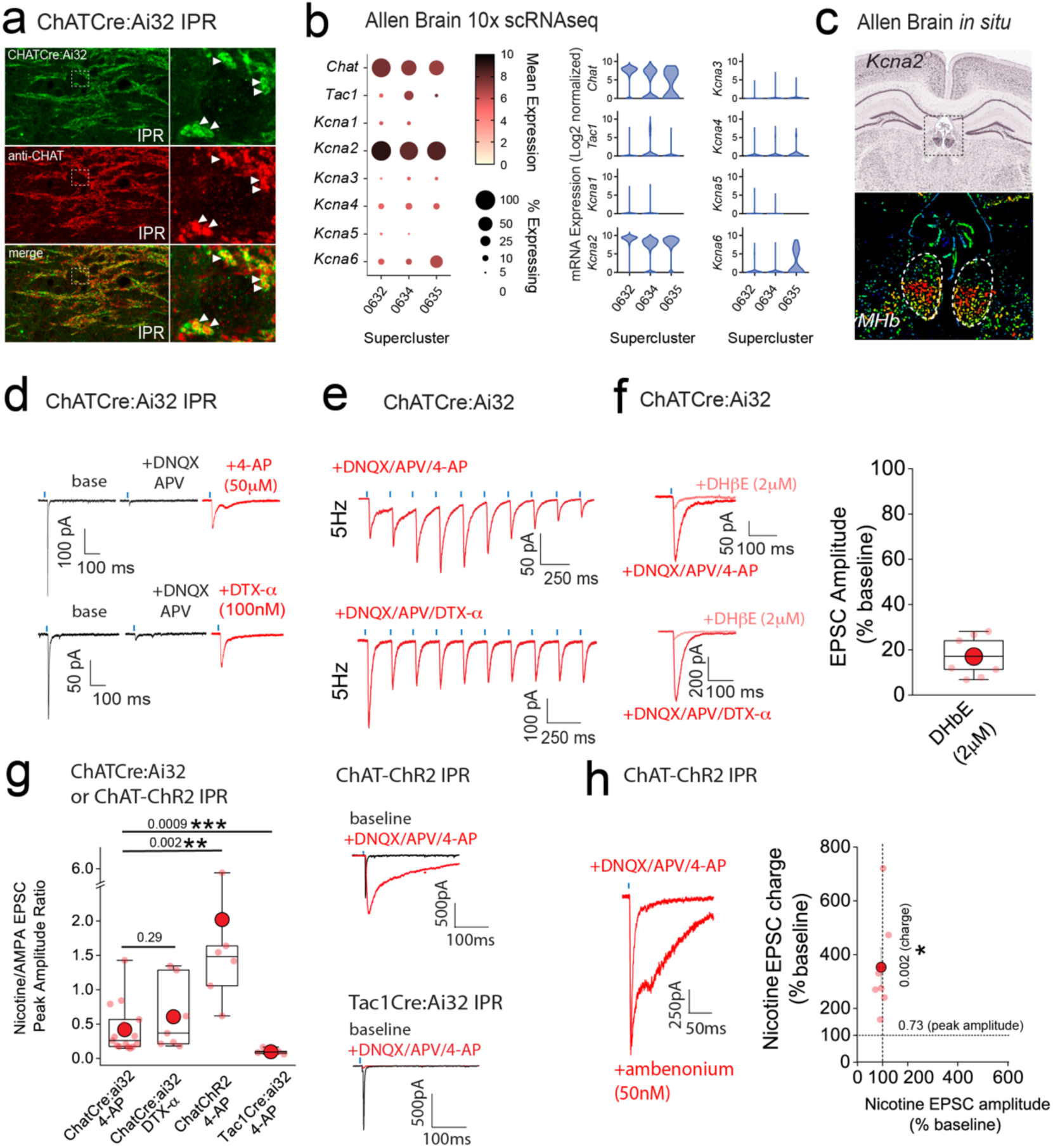
Kv1 channels constitute a molecular brake of nicotinic receptor mediated signaling in the IPN. **(a)** High resolution airy scan images of IPR in ChATCre:Ai32 mice showing ChR2-expressing cholinergic boutons (green) and endogenous ChAT (red). Right panels are magnified regions of the boxed area in left panels. Arrows indicate faithful expression of ChAT within ChatCre:Ai32 boutons (**cf.** Fig. 5a). **(b)** Corresponding dot and violin plots illustrating the relative expression of *KCNA1-6* in CHAT superclusters only. **(c)** *In situ* hybridization for *KCNA2* mRNA (top panel) and corresponding pseudo-colored expression level (bottom panel) illustrating bias towards ventral mHb. Data are from the Allen Brain Institute (https://mouse.brain-map.org/gene/show/16263). **(d)** Voltage-clamp example traces of light-evoked EPSCs mediated by cholinergic mHb neurons (left panel) under baseline, following 10μM DNQX/100μM DL-APV plus 50μM 4-AP (top panel) or plus 100nM dendrotoxin-α (bottom panel). **(e)** Voltage-clamp example traces of 5Hz trains of light-evoked EPSCs (10 stimuli) mediated by cholinergic mHb neurons in the presence of 10μM DNQX/100μM DL-APV and 100μM 4-AP (top panel) or DTX-α (bottom panel) in a ChATCre:Ai32 mouse. **(f)** Single light stimulus evoked EPSC in the presence of 10μM DNQX/100μM DL-APV and 100μM 4-AP (top left panel) or DTX-α (bottom left panel) in the absence or presence of 2μM DHβE. Box plot with individual data of the percentage inhibition of the light-driven EPSC peak amplitude mediated by cholinergic neurons in the presence of 10μM DNQX/100μM DL-APV/50μM 4-AP or DTX-α (n = 9 recorded IPR neurons from 9 mice). (**g)** Box plot of the nicotine/AMPA (nAChR/AMPA) peak amplitude ratio within individual recorded IPR neurons percentage as measured under baseline (AMPA EPSC) and in the presence of 10μM DNQX/50μM DL-APV/100μM 4-AP (nAChR EPSC; n = 15 recorded IPR neurons from 15 ChATCre:Ai32 mice) or 100nM DTX-α (nAChR EPSC; n = 7 recorded IPR neurons from 5 ChATCre:Ai32 mice). nAChR/AMPA peak EPSC amplitude ratios were also performed in ChAT-ChR2 (n = 6 recorded IPR neurons from 6 mice) and Tac1Cre:Ai32 mice (n = 6 recorded IPR neurons from 6 mice). Voltage-clamp example trace of light-evoked EPSCs under baseline and after addition of 10μM DNQX/50μM DL-APV/100μM 4-AP in ChAT-ChR2 (top right panel) and Tac1Cre:Ai32 (bottom right panel). **(h)** Voltage-clamp example trace of light-evoked nAChR-mediated EPSCs mediated by cholinergic mHb neurons under baseline and following application of 50nM ambenonium (left panel) in a ChATChR2 mouse. Scatter plot of the individual (light red symbols) and pooled (red symbol) percentage change in nAChR EPSC peak amplitude versus EPSC charge (measured over the first 500ms duration of the EPSC) in each individual recording (n = 7 recorded IPR neurons in 5 mice; right panel). Data in **(b)** and **(c)** are from the publicly available Allen Brain Cell Atlas (https://knowledge.brain-map.org/abcatlas) and the Allen Brain Map (Allen Brain; https://mouse.brain-map.org/gene/show/16263), respectively. See methods for further details.

**Figure 7-figure supplement 1.**
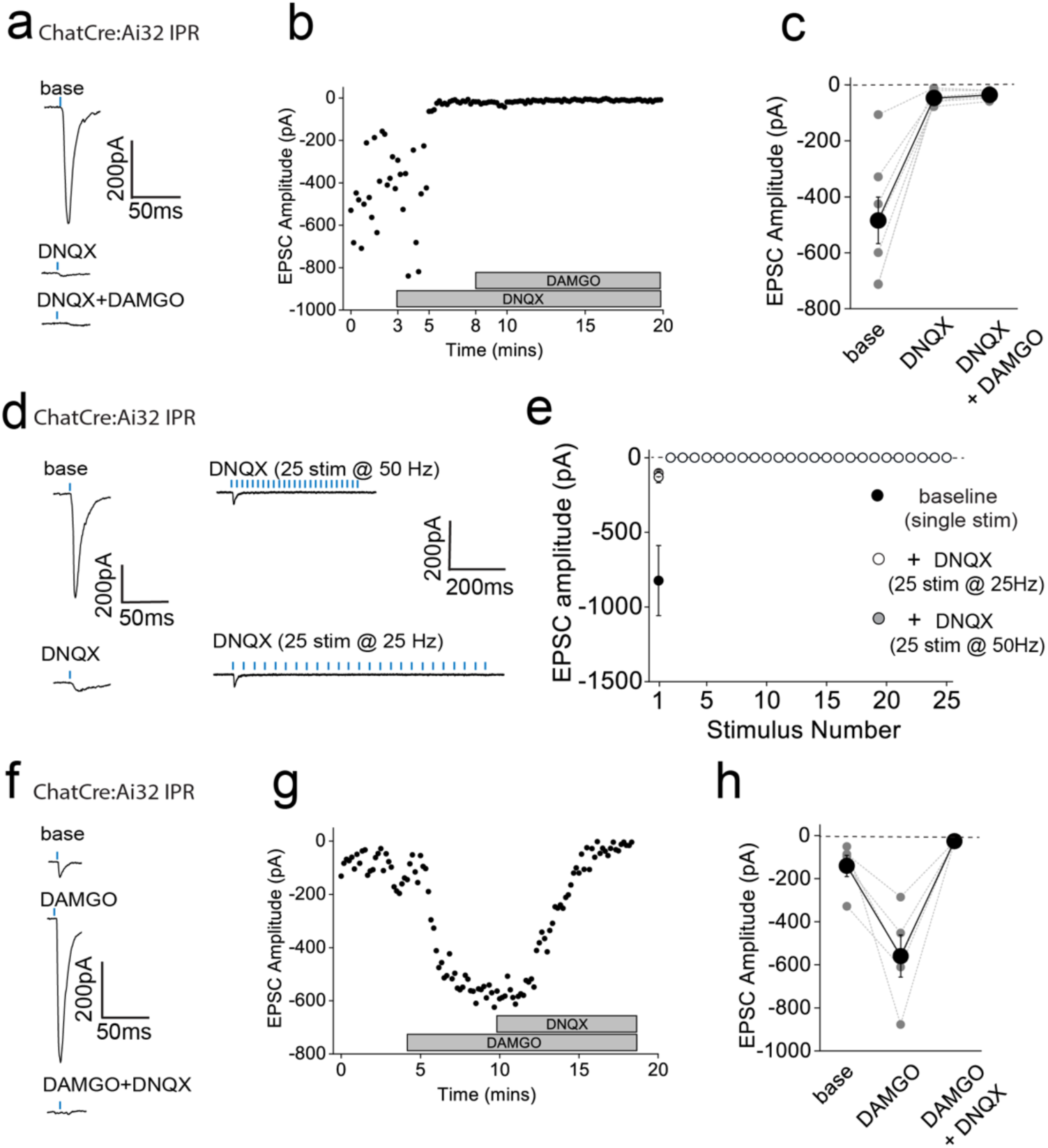
Effect on DNQX on synaptic transmission mediated by mHB cholinergic neurons onto IPN prior to or after mOR activation and during high frequency stimulation. (a,b) Voltage-clamp example traces of light-evoked AMPAR EPSCs mediated by cholinergic mHb neurons and time course of peak amplitude under baseline, 10μM DNQX and 10μm DNQX plus 500nM DAMGO conditions. **(c)** Individual (grey filled symbols) and mean (black filled symbols) data of AMPAR EPSC amplitude under baseline, 10μM DNQX and 10μm DNQX plus 500nM DAMGO conditions (n = 7 recorded IPR neurons from 5 mice). (**d**) Voltage-clamp example traces single stimulus light-evoked AMPAR EPSCs mediated by cholinergic mHb neurons stimulation in the presence and absence of 10μM DNQX (left panels). Traces from the same cell in response to trains of light (1ms, 25 stimulations given at 50 or 25Hz; right panels. (**e**) Pooled data of light evoked responses in response to single stimulation (black) and during 50 and 25 Hz trains (grey and open symbols, respectively; n = 5 recorded cells from 2 mice). **(f,g)** Voltage-clamp example trace of single light-evoked AMPAR mediated EPSCs mediated by cholinergic mHb neurons and time course of peak amplitude under baseline, 500nM DAMGO and 500nM DAMGO plus 10μm DNQX conditions. **(h)** Individual (grey filled symbols) and mean (black filled symbols) data of AMPAR EPSC amplitude under, baseline, 500nM DAMGO and 500nM DAMGO plus 10μm DNQX conditions. (n = 5 recorded IPR neurons from 5 mice).

In these experiments we were “blind” to the molecular identity of the postsynaptic IPR neuron. Since a major population of neurons in this IPN subregion are of the somatostatin (SST) GABAergic subtype^15,39^, it is likely that most of our recordings are from this subtype.

Nevertheless, we also employed ChAT-ChR2 transgenic mice containing fluorescently reported SST neurons (i.e. ChATChR2:SSTCre:Ai9; **Figure 3d**) to allow targeted recordings specifically from these neurons. Our data clearly show that DAMGO robustly increases AMPAR EPSCs impinging on IPR RFP+ SST neurons (**Figure 3d,e**) accompanied by a trending increase in PPR (**Figure 3f**). We also performed a series of additional experiments agnostic to the postsynaptic neuronal identity in IPC subdivision. Despite the relatively low expression levels of mOR in this subdivision of IPN (**Figure 1e,f)** we observed a robust increase in AMPAR EPSC amplitude at a similar extent to that seen in IPR (**Figure 3-figure supplement 1**) thus extending this remarkable potentiation of glutamatergic synaptic transmission to all IPN subfields where cholinergic neuronal terminals reside.

We considered non-specific actions of DAMGO such as those that could be mediated by a possible direct modulation of ChR2 itself to explain this non-canonical observation. However, our previous data demonstrating the reduction of glutamatergic neurotransmission mediated by SP neurons by DAMGO described (**Figure 2**) renders this possibility unlikely. Furthermore, a similar potentiation of cholinergic neuronal mediated glutamatergic transmission in the IPN upon activation of GABA_B_ receptors has demonstrated a propensity for this circuit to undergo such modulation in response to a Gi-linked receptor^32–34^. Nevertheless, despite this precedent, we employed a pharmacological approach to further validate this unexpected result. We show that lower concentrations of DAMGO (100 nM) and alternative mOR agonists, Met-enkephalin (Met-enk, 3 μM) and morphine (10 μM) all induce potentiation of ChAT neuronal glutamatergic transmission to similar extents (**Figure 3g**). In addition, pre-treatment with the mOR selective antagonist (CTAP, 1μM) completely prevents the DAMGO induced response (**Figure 3g**).

Together, the pharmacological battery of tests employing an experimental, an endogenous and clinical/recreational mOR agonists and selective mOR antagonism clearly implicate this opioid receptor in mediating a non-canonical potentiation of AMPAR EPSC amplitude elicited by mHb cholinergic neurons in IPN.

In several recordings from both “blind” patching of IPR neurons and in directly identified SST neurons, no measurable post synaptic AMPA receptor EPSC could be elicited following light-evoked stimulation of cholinergic axons even with the maximal possible light intensity available (i.e. 100 % arbitrary LED power; approximately 6.9 mW/mm^2^), and thus further interrogation was typically not performed. However, in a few instances, we nevertheless proceeded with DAMGO or Met-enk application and surprisingly, in a subset of these recordings (9/15 cells tested), a robust and reversible emergence of a significant AMPAR mediated EPSC was observed (**Figure 3h,i**). Thus, these data clearly reveal that mOR activation invokes a synaptic mechanism that, at its most extreme, results in an “un-silencing” of ChAT neuronal glutamatergic transmission in IPN including in directly identified SST GABAergic neurons.

Since, PPR measurements are only valid if the same population of release sites are assayed before and after experimental manipulation, the additional engagement of a putative reluctant vesicular pool^32^ by mOR activation could explain the discrepancy between the augmentation of EPSC amplitude and observed changes in PPR (**Figure 3b,c,e,f**). Future interrogation is warranted for definitive identification of the relative contributions of pre- and post-synaptic loci to the potentiation observed (but see^37^).

To date, we have examined the effect of mOR on synaptic transmission during paired pulse light activation delivered at 20 Hz. Although, mHb cholinergic neurons are capable of burst firing at much higher frequencies in response to afferent stimulation *in vivo*^12^, they typically exhibit low frequency (i.e. <10Hz) intrinsic (i.e. independent of synaptic input) firing *in vitro*^10,40–42^. In agreement with these previous studies, cholinergic neurons in ventral mHb (identified in ChATCre:Ai9 mice; **Figure 1c**) spontaneously elicit action potentials measured in cell-attached recordings at an average of ∼5 Hz (range 2 - 10Hz; **Figure 4a**). Therefore, we assessed the role of mOR activation on glutamatergic transmission mediated by cholinergic neurons elicited by light stimulation trains delivered at this frequency. Interestingly, even with this relatively low frequency paradigm, glutamatergic transmission is extremely labile in nature as evidenced by the large and rapid depression of AMPAR EPSC during the stimulus train (**Figure 4b,c**).

Remarkably, activation of mORs essentially eliminates activity-dependent depression at this synapse (**Figure 4b,c**). This switch in transmission dynamics greatly facilitates the probability of excitatory postsynaptic potential-spike coupling (ES coupling) in response to each stimulus within the train (**Figure 4d-e**). Thus, these data clearly demonstrate that at physiologically relevant stimulus patterns, mORs serve to dramatically increase the fidelity of glutamatergic transmission in the IPR mediated by cholinergic neurons culminating in an enhanced recruitment of postsynaptic IPR neurons.

### mORs constitute a molecular switch to alter the salience of glutamatergic transmission mediated by cholinergic and substance P neurons in the IPR

An intriguing inconsistency was observed during the conduction of our experiments regarding the DAMGO effect on light-evoked and spontaneous EPSCs (sEPSCs) impinging on IPR neurons. Under the experimental conditions employed sEPSC events were exclusively mediated by AMPARs as evidenced by their virtually complete cessation upon DNQX application (**Figure 5-figure supplement 1a**). Remarkably, in contrast to the previously described robust potentiation of light-evoked AMPAR EPSCs, activation of mORs significantly attenuates sEPSC frequency in agreement with a recent study^37^, with no effect on amplitude (**Figure 5-figure supplement 1b-d).** Thus, in a single IPR postsynaptic neuron opposing effects on light-evoked and spontaneous AMPAR EPSCs is evident (**Figure 5-figure supplement 1e**). Although these data do not directly identify the origin and neuronal subtype(s) responsible for the sEPSCs measured, the incongruent effect of mOR activation on evoked versus spontaneous events led us to consider the possible existence of an additional afferent system impinging on IPR neurons. Prevailing circuit schemas based on axonal arborization patterns portray a mutually exclusive, non-overlapping afferent input to IPN mediated by cholinergic and SP neurons to IPR/IPC versus IPL, respectively. However, in the current study by employing high-resolution imaging in Tac1Cre:Ai32 mice a clear presence of synaptic bouton-like structures in the IPR is noted (**Figure 5a,b**). These structures do not co-localize with endogenous ChAT, but do express the glutamate vesicular transporter, VGluT1 (**Figure 5a,b**).

This presence of putative presynaptic anatomical substrates points to the existence of a secondary input to IPR mediated by SP neurons distinct to the well-established one originating from mHb cholinergic neurons. Indeed, significant light-evoked AMPAR EPSCs in IPR neurons can be elicited in Tac1Cre:Ai32 mice with essentially similar amplitudes across all LED powers employed to that seen in ChATCre:Ai32 mice (**Figure 5c**). Thus, in contrast to recent ultrastructural EM analyses^32^, these data indicate a previously unidentified functional glutamatergic input to IPR mediated by SP neurons. Interestingly, DAMGO application significantly depresses the SP neuronal glutamatergic output in IPR and increases PPR (**Figure 5d-f**) to a similar extent to that observed in IPL (cf. **Figure 2**). This is in direct contrast to the role of mORs in positively modulating the ChAT neuronal glutamatergic transmission in the IPR (**Figure 3a,b,d,e**). Thus, for the first time we demonstrate both SP and cholinergic input to the same subregion of IPN and that mOR activation elicits diametrically opposing effect on transmission mediated by these respective presynaptic neuronal populations.

### Dynamic regulation of the mOR elicited afferent specific switch in salience of the glutamatergic transmission to the IPR during adolescence

Our functional studies thus far have been restricted to sexually mature, adult mice (> p40). mOR expression emerges in many brain structures during prenatal development largely overlapping with their final adult distribution profile by mid/late gestation^43^. In agreement, we reveal that mOR protein expression in IPR is present at appreciable levels from early postnatal stages (i.e. from P10 onwards; **Figure 6a**). This begs the question whether modulation of habenulo-interpeduncular synaptic transmission described during these critical early epochs mirrors that seen in the adult. To examine this, we extended our analyses to earlier time points (P15 to P40) and found that mOR modulation of glutamatergic transmission mediated by cholinergic neurons in the IPR undergoes a remarkable developmental regulation. This is characterized by an initial inhibitory effect on light-evoked AMPAR EPSCs transitioning to the previously described augmentation (**Figure 6b,c,f**). In contrast, the glutamatergic input to the IPL and the newly discovered input to the IPR elicited by SP neurons is not developmentally regulated with synaptic transmission being consistently inhibited by DAMGO application at all ages tested (**Figure 6d-f**). Thus, despite essentially similar levels of IPR mOR expression at the ages assayed (**Figure 6a**), at earlier development stages opioids result in overall inhibition of afferent input to the IPR agnostic to afferent input with the emergence of the previously highlighted differential regulation of mHB cholinergic versus substance P neuronal output emerging during the late adolescent stages (**Figure 6f).**

### mOR induced potentiation of nicotinic receptor signaling in IPR is conditional on the removal of a molecular brake mediated by Kv1 channel function

In addition to mORs, this circuitry contains dense expression of nicotinic receptors located at pre- and postsynaptic sites^10,44,45^. However, the ability to reliably evoke post-synaptic nicotinic receptor (nAChR) EPSCs in response to physiologically appropriate stimuli has been challenging. Indeed, many studies have resorted to high frequency and prolonged stimulation^36^ or bath/puff applications of nicotine to investigate nAChR function in this circuit^19,39,40,46–48^.

Despite the faithful co-expression of endogenous ChAT in ChatCre:Ai32 expressing terminals located in the IPR(**Figure 7a**), evoked transmission by brief light pulses (1-5ms) from cholinergic neurons predominantly elicits pure AMPAR EPSC in postsynaptic IPR neurons under our basal conditions as described above and in agreement with previous observations^21,36,37,39^ (**see Figure 5-figure supplement 1a and Figure 7-figure supplement 3a-e**). Additionally, DAMGO, at concentrations that strongly potentiate glutamatergic transmission, does not result in a measurable postsynaptic nAChR EPSC after AMPARs are pharmacologically silenced (**Figure 7-figure supplement 1a-c**). Furthermore, repeated light stimulation at either 25 or 50Hz fails to elicit the emergence of any nAChR mediated EPSCs in the presence of DNQX (**Figure 7-figure supplement 1d-e**). Finally, the potentiated ESPC following mOR activation is purely AMPAR mediated as evidence by complete block by DNQX (**Figure 7-figure supplement 1f-h**). Together, our functional analyses demonstrate that brief pulses of light (1-5ms) in ChatCre:Ai32 mice does not result in sufficient ACh release, if any, to elicit measurable postsynaptic nAChR responses in response to high frequency stimulus regimens and even under conditions where mOR activation strongly enhances glutamatergic transmission.

Interestingly, cholinergic transmission at the neuromuscular junction (NMJ) can be boosted via potassium channel blockade with fampridine (4-aminopyridine) and has been clinically indicated in disorders such as myasthenia gravis and multiple sclerosis^49^. Furthermore, low micromolar 4-AP in slices amplifies acetylcholine release, as assessed by Grab-ACh mediated fluorescence in the IPN^50,51^. Experimentally, these concentrations are relatively selective in inhibiting K^+^-channels of the Kv1 family. Probing the publicly available 10X single cell RNAseq database provided by the Allen Brain Institute reveals that *KCNA2* (the gene encoding Kv1.2) is the most prevalently expressed in the designated ChAT Superclusters (**Figures 1d, 7b**; mean expression levels including zero values = 9.1, 6.8 and 4.1 in Superclusters 0632, 0634 and 0635, respectively). Furthermore, spatial interrogation via *in situ* hybridization (Allen Brain; https://mouse.brain-map.org/gene/show/16263) demonstrate the presence of appreciable *KCNA2* transcript levels in ventral habenula (**Figure 7c**) a region that is enriched in cholinergic neurons (**Figure 1b,c**). We therefore speculated that Kv1.2 block could result in enhancement of ACh release to perhaps reveal postsynaptic nAChR EPSCs in IPR neurons. Indeed, following block of glutamatergic transmission (DNQX + APV), subsequent application of 4-AP (50-100 μM) or the more selective Kv1 channel antagonist, dendrotoxin-α (100 nM) results in the emergence of an EPSC elicited by brief single light pulses (**Figure 7d)** and in response to trains of 5Hz stimulation (**Figure 7e**). This emergent EPSC is mediated by nAChRs as evidenced by ∼ 80-90% block by the nicotinic receptor antagonist, DHβE^52,53^ (2 μM; **Figure 7f**).

Interestingly, this experimental approach allows for the calculation of nAChR;AMPAR peak amplitude ratio enabling an assessment of fast excitatory cholinergic transmission normalized to stimulus intensity required to produce a given AMPA EPSC response in an individual postsynaptic IPR neuron. Thus, the size of nAChR response can be directly compared across multiple recordings from differing slices, mice and experimental conditions. We first assessed the extent to which both 4-AP and DTX-α reveal light-evoked nAChR EPSCs finding no significant difference in the nAChR:AMPAR ratio, demonstrating that the specific Kv1.1/Kv1.2 channel antagonist unmasks nAChR mediated synaptic transmission in a similar manner to low concentration 4-AP (**Figure 7g)**. This, in combination with the predominant expression of *KCNA2* (**Figure 7b,c**) and relative lack of other *KCNA* transcripts (**Figure 7b**) in mHb ChAT neurons confirms our hypothesis that Kv1.2 plays a central role in unmasking nAChR-mediated synaptic transmission. In addition to ChATCre:Ai32 mice, we also employed the ChAT-ChR2 mouse. Not surprisingly, nAChR EPSCs could also be elicited in this mouse line following 4-AP application (**Figure 7g**). However, the measured nAChR:AMPAR ratio is significantly higher than that seen in ChatCre:Ai32 mice (**Figure 7g**) indicating a skew towards more prominent cholinergic transmission when compared to that mediated by glutamate. It is unclear as to the exact reason for this divergence, but it may be related to increased levels of the vesicular Ach transporter (vAChT) expression observed in ChAT:ChR2 mice^54^. Although TAC1 mHb neurons also express *KCNA2* transcript albeit lower than in the CHAT neurons (data not shown), 4-AP application following block of glutamatergic transmission does not result in emergence of any additional EPSC in response to single stimuli in Tac1Cre:Ai32 mice (**Figure 7g**). Finally, application of a cholinesterase inhibitor (ambenonium, 50 nM) markedly prolongs the EPSC waveform with minimal effect on amplitude (**Figure 7h**). Thus together, these results corroborate the role of Kv1.2 as a molecular brake, removal of which, results in the emergence of habenulo-interpeduncular synaptic transmission via postsynaptic nAChR signaling on IPR GABAergic neurons.

Having established the ability to reliably assess an otherwise reluctant cholinergic transmission, we next investigated how nAChR mediated signaling in the IPR is impacted by mOR activation. As with AMPAR mediated EPSCs, DAMGO produces a marked potentiation of light-evoked nAChR EPSC amplitude in IPR with the augmented EPSC being completely blocked by DHβE (**Figure 8a,b**). The mOR-mediated potentiation of both the glutamatergic and cholinergic output is essentially similar (**Figure 8i**). Till now, we have utilized optogenetic approaches to assay synaptic transmission in the IPN. Interestingly, following acute block of Kv1, nAChR EPSCs could also be elicited via electrical stimulation (**Figure 8c**). Further,

**Fig. 8.**
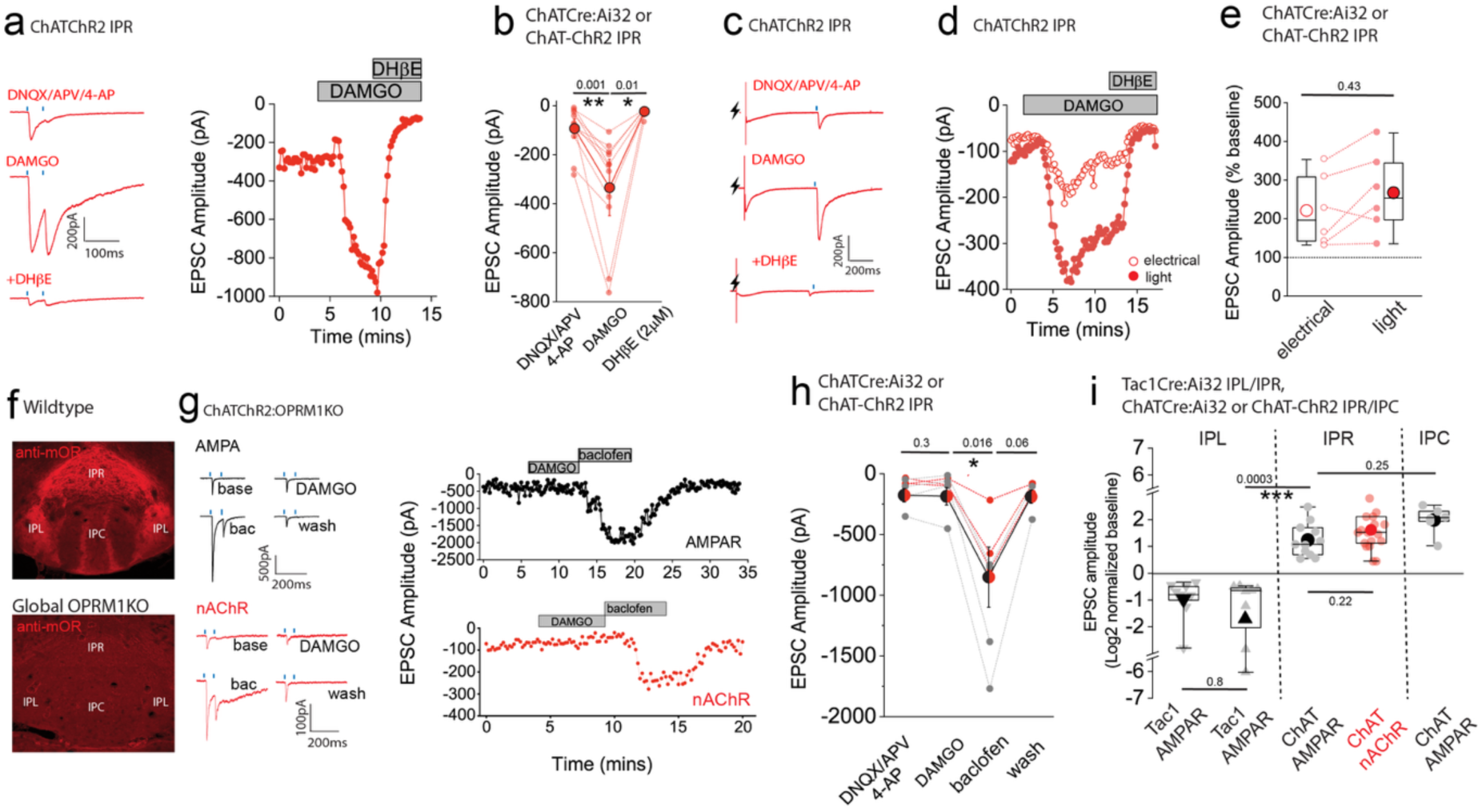
mOR potentiates nAChR EPSC amplitude revealing an interplay between opioid and cholinergic systems in the habenulo-interpeduncular axis. **(a)** Voltage-clamp example traces of light-evoked nAChR EPSCs (2 stimuli at 20Hz; blue dashes) mediated by cholinergic mHb neurons (left panel) and time course of peak amplitude (right panel) under baseline, 500nM DAMGO and 500nM DAMGO plus 2μm DHβE conditions. **(b)** Individual and mean (red filled symbols) data of nAChR EPSC peak amplitude. **(c,d)** Voltage-clamp example traces of simultaneous electrical (lightning symbol) and light-evoked (blue dash) nAChR EPSCs and time course of peak amplitude (open and filled symbols representing electrical and light-evoked peak amplitude of nAChR EPSCs, respectively) under baseline, 500nM DAMGO and 500nM DAMGO plus 2μm DHβE conditions. **(e)** Percentage change in electrical and light-evoked nAChR EPSC peak amplitude elicited by 500nM DAMGO in each individual recorded IPR neuron (n = 6 recorded neurons from 6 mice). **(f)** Confocal images of endogenous mOR protein expression in IPN of WT and homozygote OPRM1 KO mice. **(g)** Voltage-clamp example traces of light-evoked AMPA and nAChR EPSCs (2 stimuli at 20Hz; blue dashes) mediated by cholinergic mHb neurons (top and bottom left panels, respectively) and time course of peak amplitude (right panels) under baseline, 500nM DAMGO, 1μM baclofen and washout conditions in ChATChR2:OPRM1KO mice. **(h)** Individual and mean data of light-evoked AMPAR (black symbols; n=4 from 4 mice) and nAChR (red symbols; n=3 from 3 mice) peak amplitude in response to DAMGO and baclofen application in ChATChR2:OPRM1KO mice. Note in 2 of the 4 AMPAR EPSC recordings washout of the baclofen effect was not performed. **(i)** Summary box plot with individual data of the percentage normalized changes (log2) of AMPAR and nAChR peak amplitudes in response to mOR activation mediated by SP and cholinergic neurons in various subdivisions of IPN tested. Note that the normalized data in this panel are replotted from the absolute peak EPSC amplitude changes mediated by mOR agonists in **Figures 2b, 3b,3f, 5e**, **7b** and **Figure 3-figure supplement 1b.**

DAMGO reliably potentiated both electrical and light evoked nAChR EPSCs in a single IPR neuron to similar extents, a response that is effectively blocked by DHβE (**Figure 8c-e**). Finally, we tested the molecular specificity of the DAMGO response employing ChAT-ChR2:global *OPRM1* knockout mice. In these mice DAMGO application is ineffective in augmenting glutamatergic or cholinergic mediated neurotransmission in IPR whilst the potentiation upon GABA_B_ receptor activation previously described^32,34,55^ remains intact (**Figure 8f-h**; note that in these experiments CGP55845A which is routinely included in all other experiments was omitted). This genetic approach complements our previous pharmacological data (**Figure 3g**) to directly implicate mORs in the potentiation of mHb-IPN transmission. The overall effects of mOR activation on synaptic transmission mediated by the distinct mHb afferent systems in the various subdivisions of IPN tested is summarized in **Figure 8i**.

## DISCUSSION

In the current study, we reveal a remarkable augmentation of glutamatergic/cholinergic co-transmission by mHb cholinergic neurons following mOR activation in the IPN. However, the underlying cellular/network mechanisms responsible remain unclear. The most parsimonious explanation is that mORs in proximity to the release machinery of cholinergic nerve terminals^23^ mediate this effect. However, one must consolidate the fact that a Gi-linked receptor, which typically serve to directly inhibit release, results in a seemingly paradoxical potentiation.

Functional and mechanistic studies have elegantly demonstrated that GABA_B_ receptors (another member of the Gi-linked subfamily) trigger acute molecular and structural adaptations resulting in enhanced Ca^2+^ influx and a switch of transmission modes within single presynaptic terminals to ultimately increase neurotransmitter release^32,33^. Although, these observations set a precedent for potentiation by Gi-linked receptor activation in the IPN similar experimental approaches to that employed in the examination of the GABA_B_-receptor mediated potentiation^32,33,51^ are required to definitively implicate overlapping mechanisms. In other brain regions, a canonical inhibitory influence of mORs elicits overall network excitation via disinhibition^56–59^. Furthermore, numerous studies have revealed an intricate interplay involving diverse neuromodulatory components intrinsic to the IPN that serve to regulate synaptic transmission^15,55,60^. Thus, taking the results of the current study in isolation, we cannot discount the possibility that the potentiation observed here may result via the activation of mORs residing on other neural elements within the IPN microcircuitry such as postsynaptic neurons or glia, for instance. However, it must be noted that a recent study employing a viral strategy to elicit conditional knockout of mORs in *OPRM1*-expressing mHB neurons specifically, prevents the

DAMGO mediated potentiation of glutamatergic transmission onto the IPN indicating a presynaptic locus^37^. Regardless, of the exact underlying cellular and/or network mechanisms our data extend the previously described Gi-linked receptor potentiation of neurotransmission in the IPN^32,33,51^ to mORs, a predominant target of a societally relevant and prevalently misused class of drugs. It would be of interest to determine whether GABA_B_ receptor activation exerts a similar inhibitory influence as mORs on the newly discovered SP neuronal mediated transmission to the IPR. In addition, does the modulation of ChAT neuronal glutamatergic output by GABA_B_ receptors undergo a similar developmental regulation to that observed with mORs?

These additional functional comparisons of the synaptic influences of these two distinct Gi-linked receptors may shed light as to the similarity, or lack thereof, regarding the respective underlying cellular mechanisms.

Recent generation of transgenic mice^24,61,62^ have permitted manipulation of mOR-expressing neurons and mOR receptors including those expressed specifically within the mHb/IPN. Emerging behavioral studies adopting such conditional genetic approaches have highlighted an important role of mORs within this circuitry in mediating varying aspects of reward and aversion^25,26^. Unsurprisingly, it has been concluded that the observed behavioral effects of such manipulations are due to perturbations of a generalized inhibition mediated by mORs within the mHb/IPN circuit. Here, we reveal a functional dichotomy characterized by an inhibitory yet excitatory influence on neurotransmission mediated by SP and cholinergic neurons, respectively. These distinct neuronal populations participate in and drive specific behavioral facets demonstrating a division of labor. For example, SP neuronal activity positively correlates with reward outcome, history and hedonic value^63,64^, whereas that of cholinergic neurons is reduced during behaviors associated with reward^63^. mHb cholinergic neurons have also been linked to negative affect including anxiety and depression^42,65,66^ that promote drug-seeking behaviors during withdrawal^67^. We reveal for the first time that the IPR subdivision receives functional glutamatergic afferent input from SP neurons that based on previous anatomical interrogation were considered to exclusively target the IPL^68^. Interestingly, the novel opposing effect on transmission extends to the IPR thus demonstrating a role for mORs in altering the relative salience of these distinct afferent systems with regard recruitment of common downstream IPR neurons. The IPR houses somatostatin (SST) GABAergic neurons whose afferents impinge on the raphe nucleus and lateral dorsal tegmental nucleus, the latter influencing the VTA and, in turn, its downstream structures^15,39,69^. Thus, careful consideration regarding future examination of behavioral outcomes resulting from the described contrasting effects of mOR activation on the parallel processing of reward (mHB SP) and anti-reward (mHB ChAT) in this circuit is essential.

A limitation of the current study arises from the sole utilization of a transgenic approach to selectively assay light-evoked synaptic transmission from ChAT and SP neuronal populations, respectively. Although the mHb provides most of the afferent input to the IPN, this approach does not exclude possible activation of additional inputs from other brain regions^22,70,71^. This caveat can be circumvented by use of stereotaxic viral delivery to express ChR2 solely in the mHB^21,37,39^. However, it is unclear if neuronal inputs from these possible alternate sources^22,70,71^ are glutamatergic in nature and mediated by a TAC1/OPRM1-expressing neuronal population.

Nevertheless, to definitively identify the described novel input as one that originates from mHb SP neurons will require the future use of such viral strategies.

Throughout this study we contextualize the effect of mOR activation in the IPN primarily through the lens of exogenously introduced opioids. However, physiological activation of mORs can be mediated to differing extents by endogenous ligands such as dynorphin or enkephalin. Indeed, we demonstrate that met-enkephalin effectively boosts co-transmission from cholinergic neurons in a similar manner to morphine. Interestingly the IPN is home to a population of pro-enkephalin (PENK)-expressing neurons and detectable met-enkephalin immunoreactivity^23,72^. In other brain regions, manipulation of PENK neuronal activity and or ablation of PENK itself elicits mOR mediated circuit modulation and behavioral alterations^73–75^. It is anticipated that activation of mORs by local endogenous opioid release yields similar complex effects to those following exogenous application of mOR agonists described here. Thus, this physiological route will have major implications concerning the role of mHB/IPN mORs in mediating the positive reinforcing effects of non-opioid drugs of abuse (e.g. nicotine, alcohol, amphetamines) and other “natural” rewarding stimuli (e.g. such as those associated with social interaction, exercise and food intake) thus extending the relevance of our study to these additional modalities.

Another novel finding in the current study relates to the identification of a “molecular brake” that exerts strong control over nAChR signaling impinging on postsynaptic IPN neurons. Specifically, compromising the function of delayed rectifying K^+^-channels containing Kv1.2 subunits results in emergence of nAChR EPSCs in response to light and electrical stimuli delivered with physiological paradigms likely thru increased axonal/bouton excitability and calcium influx via action potential waveform modulation. Kv1.2 function can be bidirectionally impacted by use-dependent mechanisms or through secondary activity mediated cascades resulting in post-translational modifications (e.g. phosphorylation state) that impact its function and/or trafficking^76–78^. These plausible endogenous mechanistic routes could serve to titer the strength nAChR mediated transmission in the IPR.

The resident nAChRs in the mHb/IPN are primarily encoded by the CHRNA3/B4/A5 gene cluster and the role of this circuitry in nicotine use is well characterized. For example, increased propensity for nicotine abuse in humans is associated with dysregulation of nAChR function precipitated by single nucleotide polymorphisms of this gene cluster^79–81^. Experimental manipulation of nAChR signaling in the mHb/IPN generates phenotypes associated with various aspects of nicotine consumption in mouse models^53,82–85^. Furthermore, chronic nicotine results in adaptations in nAChR expression and function that can exacerbate continued nicotine consumption^40,48,86,87^. Together, these studies establish a link between nAChR mediated signaling within this circuitry and prevalence of nicotine misuse. Here we highlight an intriguing interplay between opioid and cholinergic systems within the mHb/IPN axes. Our data clearly demonstrate that mORs and Kv1.2 together comprise synergistic molecular targets that, in addition to others previously identified^34^, when leveraged in tandem can manipulate nAChR mediated signaling to yield potential interventions for nicotine overuse.

Adolescence is a critical period for many aspects of brain and social development and represents a high-risk demographic group for drug use developing into long-term addiction^88^. Propelled by the societal introduction of highly potent synthetic morphine analogs (e.g. fentanyl), one of the many devastating sequalae of the well-documented opioid crisis is an alarming high rate of overdose deaths not only in adults but also during vulnerable teenage years. Of note is the striking valence conversion from depression to potentiation of mHb ChAT neuronal signaling in the IPR occurring around the late postnatal stages. This is in stark contrast to that seen with the newly described SP neuronal mediated transmission in this same IPN subdivision where mOR activation is inhibitory at all life stages assayed. Thus, during postnatal development opioids acting via mORs result in a blanket inhibition of these two distinct inputs prior to the establishment of the afferent specific modulation in the adult. Interestingly, numerous studies have highlighted changes in the role of mORs in reward associated behaviors coinciding with similar periods^89,90^. An attractive hypothesis is that coordinated modifications at the cellular, molecular and/or network level underlie this dynamic developmental regulation. Thus, our circuit analyses provide insights into the neural correlates for the differing propensity of mORs to regulate not only positive reinforcement but also the negative effects associated with withdrawal at various stages of development^91^. Whether these adaptations underlie the increased propensity for substance abuse during development remains to be ascertained. Furthermore, opioid exposure during early developmental epochs imparts long lasting effects on the brains reward circuitry^3^. Thus together, it is imperative to consider the newly described developmentally regulated impact of mORs in the modulation of the mHb/IPN circuitry. This is of particularly relevance with respect to generation of potential preventative treatments for the deleterious consequences of fetal opioid exposure and SUDs in high-risk juveniles.

In addition to a central role in addiction, the mHb/IPN circuitry encodes fear associated behaviors^32,33,92–94^. Interestingly, prevention of GABA_B_-receptor function in mHb cholinergic neurons or increasing IPN neuronal activity facilitates and augments expression of fear extinction, respectively^32,33^. Dysregulation of this behavioral aspect is a key underlying cause of post-traumatic stress disorder (PTSD)^95^. Based on this previous work, the novel mOR mediated potentiation of the mHb cholinergic transmission and hence IPN neuronal recruitment described here constitutes a parallel, yet alternative circuit mechanism involved in the extinction of learnt fear. Thus, pharmacogenetic manipulation of mOR mediated signaling specifically in the IPN may constitute a viable approach to alleviate conditions associated with abnormal fear processing such as those seen in PTSD.

In summary, we highlight several previously undescribed and hence unappreciated roles of mORs in the dynamic regulation of synaptic transmission impinging on IPN GABAergic neurons following exposure to exogenous opioids. Although additional studies are needed to reveal the underlying cellular mechanisms responsible for these complex, divergent facets of opioid mediated effects in the habenulo-interpeduncular axis, the current detailed functional analyses lay out a valuable roadmap. One that necessitates consideration in future interrogation concerning the role of these receptors in this relatively understudied brain circuitry that encodes aspects of emotion, hedonic and addiction behaviors.

## METHODS

### Animals

All experiments were conducted under an active animal protocol approved by the Animal Care Use Committee of the National Institute of Child Health and Human Development. All transgenic mice were attained from The Jackson Laboratory (ME, USA) and were as follows: ChATCre (strain #031661), Tac1Cre (strain #021877), SSTCre (strain #018973); Ai9 (strain #007909), Ai32 (strain #024109), ChAT-ChR2-EYFP (strain #014546); OPRM1 KO (strain #007599). For conditional expression of td-Tomato (Ai9) or ChR2 (Ai32) both male and female breeders were homozygous, and all immunocytochemical or electrophysiological experiments were performed in their progeny that were heterozygous for both the Cre and floxed alleles. Heterozygous ChAT-ChR2 mice were used throughout the study either alone or crossed with the OPRM1KO mouse (homozygous) or SSTCre:Ai9 (heterozygous for both alleles). Both male and female mice were investigated, and the data were pooled.

### Reagents

Pharmacological reagents used in this study are as follows: DNQX (Cat. No. 2312/10), DL-APV (Cat. No 0105/10), CGP55845A (Cat. No. 1248/10), Picrotoxin (Cat. No. 1128), bicuculline (Cat. No. 0109/10), DAMGO (Cat. No. 1171/1), CTAP (Cat. No. 1560/1) and DHβE (Cat. No. 2349) were all purchased from Biotechne/Tocris (MN, USA); Dendrotoxin-α (Cat No. D-350 ) was purchased from Alomone Labs (Jerusalem, Israel). met-enkephalin (Cat. No. M6638) was purchased from MilliporeSigma (MA, USA). Ambenonium (Cat. No. sc-203507) was purchased from Santa Cruz Biotechnology (TX, USA). Morphine Sulphate (Cat. No. NDC: 0641-6125) was from Hikma (NJ, USA) and attained via the NIH Division of Veterinary Resources (DVR) Pharmacy.

Primary and corresponding secondary antibodies used in this study for immunocytochemistry and the working dilutions employed are as follows: Guinea pig anti-RFP (Synaptic Systems, Cat. No. 390005, 1:500) with CF555-conjugated Donkey anti-Guinea Pig IgG (Biotium Cat. No. 20276, 1:1000).Chicken anti-GFP (AvesLabs, Cat.No. GFP-1010, 1:1000) with CF488-conjugated Donkey anti-Chicken IgY (Biotium Cat. No. 20166, 1:1000 working dilution). Rabbit anti-MOR (ABCAM, Cat. No. ab134054, 1:500) with CF555-conjugated Donkey anti-Rabbit IgG (Biotium cat#20038, 1:1000). Goat anti-ChAT, Millipore (Cat. No. AB144P, 1:1000) with CF555-conjugated Donkey anti-Goat IgG (Biotium, Cat. No. 20039, 1:1000 working dilution).

### Immunocytochemistry and Imaging

Mice (P20 and older) were perfused trans-cardinally using 4% PFA and dissected brain tissues were post-fixed in 4% PFA for 24 hours at 4 °C. For P10 mice, brains were removed and drop fixed in 4% PFA for 24 hours at 4 °C. P10 mouse brains were not perfused but brains removed and drop-fixed as stated above. Fixed brain tissues were thoroughly washed in 1x phosphate buffer (PB) followed by cryopreservation using 30% sucrose. 50 μm coronal sections were made on a frozen microtome. To perform floating section IHC, brain slices were washed with 1x PB at room temperature for 1 hour with 2-3 changes of 1x PB followed by blocking and permeabilizing in Blocking Solution (1x PB + 10% donkey serum + 0.5% Triton X-100) at room temperature for at least 2 hours. Blocked brain slices were incubated in primary antibodies, which were diluted to working concentration using Antibody Solution (1x PB + 1% donkey serum + 0.1% Triton X-100), at 4 °C for 24-48 hours. After wash with 1x PB at room temperature for 15 minutes with 3 repeats, brain slices were incubated in secondary antibodies diluted with Antibody Solution at room temperature for 1 hour, followed by DAPI staining at 1ug/ml for additional 15 minutes. After washed with 1x PB at room temperature for 15 minutes with 3 repeats, brain slices were mounted on gelatin coated slides followed by air drying, covered with No.1.5 cover slides ProLong Glass Antifade Mountant (ThermoFisher Scientific, Cat#P36984), cured in darkness overnight before imaging. Confocal images were acquired on a Zeiss LSM900 using ZenBlue software. 10x and 20x confocal images of whole habenula and IPN were acquired as tiles with z-steps of 3μmx5 and 0.9μmx20, respectively. Airy scan images were acquired at 63x using multiplex 2Y mode to achieve super resolution. 2μm depth of images were captured at z-steps of 0.2μmx10, with or without tiling. All comparable images of tissues with different genotypes or developmental stages were acquired using the same light path configuration, channel input, gain and offset. Images were imported into Fiji and Adobe Photoshop for processing and densitometry measurements where applicable.

### Electrophysiology

P15-P91 mice (for details refer to the results section) were anesthetized with isoflurane, and the brain removed in ice–cold partial sucrose substituted ASCF (ssACSF) solution containing (in mM): 90 Sucrose, 80 NaCl, 25 NaHCO3, 1.25 NaH2PO4, 3.5 KCl, 4.5 MgCl2, 0.5 CaCl2, 10 glucose, saturated with 95% O_2_ and 5% CO_2_ (pH 7.4; osmolarity 300-310 mOsm). Coronal sections containing wither the mHb or IPN (270 - 300 µm) were cut using a VT–1000S vibratome (Leica Microsystems, Bannockburn, IL) in ice-cold ssACSF. The slices were allowed to recover in the ssACSF at 31-33°C for 20-30 minutes followed by an additional 20 minutes at room temperature. Slices were then transferred to our standard extracellular solution (ACSF) of the following composition (in mM): 130 NaCl, 24 NaHCO_3_, 3.5 KCl, 1.25 NaH_2_PO_4_, 2.5 CaCl_2_, 1.5 MgCl_2_, and 10 glucose, saturated with 95% O_2_ and 5% CO_2_ (pH 7.4; osmolarity 300-310 mOsm) for storage until electrophysiological recording. Slices were transferred to an upright Olympus BX51WI microscope and visualized with infrared differential interference contrast microscopy and perfused (2-4 ml/min) with the above extracellular solution. Unless otherwise indicated the recording extracellular solution was routinely supplemented with 2μM CGP55845A hydrochloride (Biotechne/Tocris Cat#1248), 50μM picrotoxin (Biotechne/Tocris Cat#1128) and 10 μM bicuculline methobromide (Biotechne/Tocris Cat#0131).

All recordings were performed at 31–33°C with electrodes (3–5 MΏ) pulled from borosilicate glass (World Precision Instruments, Sarasota, FL) filled with one of two intracellular solutions (in mM); (i) 135 CsMeSO4, 8 KCl, 4 MgATP, 0.3 NaGTP, 5 QX–314, 0.5 EGTA and 3 mg/ml biocytin. (ii) 150 K–gluconate, 3 MgCl_2_, 0.5 EGTA, 2 MgATP, 0.3 Na2GTP, 10 HEPES and 3 mg/ml biocytin. pH was adjusted to 7.3 with KOH or CsOH and osmolarity adjusted to 285–295 mOsm. Whole–cell patch–clamp recordings were made using a Multiclamp 700B amplifier (Molecular Devices, Sunnyvale, CA). Signals were filtered at 4 – 10 kHz and digitized at 10 – 20 kHz (Digidata 1322A and pClamp 10 Software; Molecular Devices, Sunnyvale, CA). For cell attached recordings pipettes were filled with our standard extracellular solution.

For optogenetic stimulation of ChR2 light stimuli at a wavelength of 470nm and duration of 1-5 ms were delivered through the 40X water immersion objective using a CoolLED pE–4000 Illumination system (Andover, UK). Arbitrary LED power typically ranged between 10 – 100% corresponding to ∼ 0.4 – 6.9 mW/mm^2^ as measured with a digital power meter (Thor Labs, NJ, USA). Where applicable, electrical stimulation (0.1 ms; typically, 100-700 μA) were performed by placement of a tungsten bipolar electrode in the IPR (World Precision Instruments FL, USA; Catalog # TST33A05KT) and a constant current stimulus isolator (World Precision Instruments; Catalog # A385 or A365). sEPSC analyses was performed on post hoc filtered episodic traces (Clampfit; Bessel 8-pole, -3db cutoff 1000Hz). Frequency of sEPSCs and average sEPSC amplitude were determined over 5 consecutive sweeps totalling a period of 4 seconds (i.e. 0.8s per sweep). sEPSC were detected via a thresholding procedure set at 4 times the RMS baseline noise. Note that only neurons with a baseline sEPSC frequency of > 8Hz were selected for further interrogation. Analyses of electrophysiological data was performed in Clampfit (Molecular Devices, Sunnyvale, CA) and curated in Microsoft Excel spreadsheets.

### Analyses of publicly available single-cell RNA sequencing transcriptomic data

We utilized the Allen Institute dataset of the single-cell transcriptomes of ∼4 million cells across the entire mouse brain^35^. To probe the region and genes of interest, we adapted Python notebooks provided for accessing the Allen Brain Cell (ABC) Atlas (https://alleninstitute.github.io/abc_atlas_access/intro.html). The log2-normalized expression matrices were downloaded and filtered to reach the cell types of interest within the habenula class “17 MH-LH Glut” (∼10.8k cells). For the current study we focused our analyses on the medial habenula subclass “145 MH Tac2 Glut” (∼8k cells). Within this subclass, we probed the 5 supertypes “0632-0636 MH Tac2 Glut”, querying the genes of interest: *Chat*, *Tac1*, *Slc17a7*, *Kcna1-6* and *Oprm1*. *Chat* supertypes were considered those with mean *Chat* expression including zero values > 3 (0632, 0634, and 0635; 6803 cells), while Tac1 supertypes were defined as those with mean *Tac1* expression including zero values > 3 (0633; 878 cells). Violin and dot plots depicting gene expression levels were created in Graphpad Prism.

### Statistical Analyses

Tests for normality (Shapiro-Wilk test) were performed on all datasets to be compared prior to statistical analyses and appropriate parametric (unpaired or paired t-tests) or non-parametric (Mann-Whitney U or Wilcoxon signed rank tests) were accordingly performed. Exact p-values are stated within the figures and values of p<0.05, <0.01 or p<0.001 were deemed statistically significant and designated with 1,2 or 3 asterisks, respectively. Error bars in the figures represent standard error of the means unless otherwise indicated. Box whisker plots were constructed as follows: symbol denotes mean value; line represents the median value; lower and upper box limits represent 25^th^ and 75^th^ percentiles, respectively. Lower and upper whiskers represent minimal and maximal data points, respectively.

## AUTHOR CONTRIBUTONS

RC was responsible for project conceptualization, experimental design, electrophysiology experiments, image processing, manuscript writing and figure preparation. CMcB, RC, KAP supervised the project. XY, AV, EL and SH conducted IHC and imaging. AC constructed modified Python notebooks provided by the Allen Brain Institute to enable analyses and representation of their publicly available mouse whole brain scRNA sequencing data. DA and SH provided critical support for mouse breeding, genotyping and colony maintenance.

## ACKNOWLEDGMENTS

We would like to thank Dr. Cole Malloy for critical discussion throughout the conduction of the study. This work was supported by an NICHD Intramural Research Program (IRP) grant to CJM (1ZIAHD001205-32). This research was supported by the Intramural Research Program of the National Institutes of Health (NIH). The contributions of the NIH author(s) are considered Works of the United States Government. The findings and conclusions presented in this paper are those of the author(s) and do not necessarily reflect the views of the NIH or the U.S. Department of Health and Human Services.

## DATA AVAILABILITY

All analyses of data associated with the manuscript have been made available: https://data.mendeley.com/datasets/3dtkmss9xz/2. Adapted Python notebooks employed for analyses and graphical representation of publicly available Allen Brain Cell Atlas mouse whole brain scRNA sequencing data can be accessed via the following link: https://github.com/acaccavano/ABC-Atlas_Chittajallu2025/.

